# Single cell transcriptome analysis defines heterogeneity of the murine pancreatic ductal tree

**DOI:** 10.1101/2020.10.12.336784

**Authors:** Audrey M. Hendley, Arjun A. Rao, Laura Leonhardt, Sudipta Ashe, Jennifer A. Smith, Simone Giacometti, Xianlu L Peng, Honglin Jiang, David I. Berrios, Mathias Pawlak, Lucia Y. Li, Jonghyun Lee, Eric A. Collisson, Mark Anderson, Gabriela K. Fragiadakis, Jen Jen Yeh, Jimmie Ye Chun, Grace E. Kim, Valerie M. Weaver, Matthias Hebrok

**Affiliations:** Diabetes Center, University of California, San Francisco, San Francisco, California, USA; CoLabs, University of California, San Francisco, California, USA; Bakar ImmunoX Initiative, University of California, San Francisco, California, USA; Department of Pharmacology, University of North Carolina at Chapel Hill, Chapel Hill, North Carolina, USA; Lineberger Comprehensive Cancer Center, University of North Carolina at Chapel Hill, Chapel Hill, North Carolina, USA; Division of Hematology and Oncology, Department of Medicine and Helen Diller Family Comprehensive Cancer Center, UCSF, San Francisco, California, USA; Evergrande Center for Immunologic Diseases, Harvard Medical School and Brigham and Women’s Hospital, Boston, Massachusetts, USA; Department of Medicine, Division of Rheumatology, University of California, San Francisco, California, USA; Department of Surgery, University of North Carolina at Chapel Hill, Chapel Hill, North Carolina, USA; Parker Institute for Cancer Immunotherapy, San Francisco, California, USA; Department of Pathology, University of California, San Francisco, San Francisco, California, USA; Center for Bioengineering and Tissue Regeneration, UCSF, San Francisco, California, USA

**Author notes:** These authors contributed equally (Arjun A. Rao, Laura Leonhardt, Sudipta Ashe).

**Keywords:** scRNA-seq, *Gmnn*, *Spp1*, pancreatic duct ligation, duct heterogeneity

## Abstract

Lineage tracing using genetically engineered mouse models is an essential tool for investigating cell-fate decisions of progenitor cells and biology of mature cell types, with relevance to physiology and disease progression. To study disease development, an inventory of an organ’s cell types and understanding of physiologic function is paramount. Here, we performed single-cell RNA sequencing to examine heterogeneity of murine pancreatic duct cells, pancreatobiliary cells, and intrapancreatic bile duct cells. We describe an epithelial-mesenchymal transitory axis in our three pancreatic duct subpopulations and identify SPP1 as a regulator of this fate decision as well as human duct cell de-differentiation. Our results further identify functional heterogeneity within pancreatic duct subpopulations by elucidating a role for Geminin in accumulation of DNA damage in the setting of chronic pancreatitis. Our findings implicate diverse functional roles for subpopulations of pancreatic duct cells in maintenance of duct cell identity and disease progression and establish a comprehensive road map of murine pancreatic duct cell, pancreatobiliary cell, and intrapancreatic bile duct cell homeostasis.

**SIGNIFICANCE:** Murine models are extensively used for pancreatic lineage tracing experiments and investigation of pancreatic disease progression. Here, we describe the transcriptome of murine pancreatic duct cells, intrapancreatic bile duct cells, and pancreatobiliary cells at single cell resolution. Our analysis defines novel heterogeneity within the pancreatic ductal tree and supports the paradigm that more than one population of pancreatic duct cells harbors progenitor capacity. We identify and validate unique functional properties of subpopulations of pancreatic duct cells including an epithelial-mesenchymal transcriptomic axis and roles in chronic pancreatic inflammation.

## INTRODUCTION

Pancreatic duct cells, while a minority of the composition of the pancreas, play an integral role in secretion and transport of digestive fluid containing proenzymes synthesized by acinar cells, electrolytes, mucins, and bicarbonate. They can serve as a cell of origin for pancreatic ductal adenocarcinoma (PDA) (1, 2) and have been implicated in the pathophysiology of multiple other diseases including cystic fibrosis (3) and pancreatitis (4).

Heterogeneity of a cell type becomes increasingly important in the context of disease and regeneration since different subpopulations can be the driving forces behind pathogenesis. The function of exocrine pancreatic cells is required for survival, yet these cells exhibit limited regenerative capabilities in response to injury. Chronic pancreatitis (CP) is a risk factor for pancreatic cancer. The underlying mechanisms for PDA progression in CP patients are incompletely understood and are likely multifactorial, including both genetic and environmental insults (5). Studies have shown that cytokines and reactive oxygen species generated during chronic inflammation can cause DNA damage. It has been hypothesized that an unlucky pancreatic cell might acquire DNA damage in the protooncogene *KRAS* or tumor suppressor genes *TP53* or *CDKN2A*, thereby accelerating malignant transformation (6, 7). Thus, it is imperative to understand the mechanisms by which DNA damage occurs in the setting of CP. Duct obstruction is one cause of CP, and the ability of ductal cells to acquire DNA damage in the setting of CP is incompletely understood.

In this report, we conducted single-cell RNA sequencing (scRNA-seq) on homeostatic murine pancreatic duct, intrapancreatic bile duct, and pancreatobiliary cells using a DBA^+^ lectin sorting strategy, and present a high-resolution atlas of these murine duct cells. By extensively comparing our subpopulations to previously reported mouse and human pancreatic duct subpopulations (8–10), we both corroborate several previous findings and identify and validate novel duct cell heterogeneity with unique functional properties including roles for subpopulation markers in CP. Our findings suggest that multiple duct subpopulations retain progenitor capacity, which is influenced by expression of markers driving subpopulation identity.

## RESULTS

### scRNA-seq identifies multiple pancreas cell types with DBA lectin sorting

Previously reported subpopulations of murine pancreatic duct cells were identified by single cell analysis of pancreatic cells obtained using an islet isolation procedure; thus, exocrine duct cells were of low abundance (9). To circumvent this issue, we employed a DBA lectin sorting strategy that has been extensively used to isolate and characterize all murine pancreatic duct cell types (11, 12), to investigate murine duct heterogeneity. We isolated live DBA^+^ cells from the pancreata of four adult female C57BL/6J littermates, and performed scRNA-seq on the pooled cells using the 10X Genomics platform (Figure 1A and S1A). After filtering out doublets and low-quality cells (defined by low transcript counts), our dataset contained 6813 cells. Clustering analysis identified 16 distinct cell populations with an average of 5345 transcripts per cell and 1908 genes per cell (Figure 1B and Dataset S1). Significantly differentially expressed genes (DEGs) when comparing a cluster to all other clusters are listed in Dataset S2. Annotation of these 16 clusters was accomplished by analysis of known markers (Figure 1B-D). Our dataset comprises 2 populations of ductal cells, a cluster of endothelial cells, one cluster of fibroblasts, and 12 immune cell clusters. As expected, murine endocrine and acinar cells are not present in our dataset because they are not DBA^+^ cells. Gene and transcript counts for each cluster are shown in Figure S1B. We identified DBA^+^Collagen I^+^ fibroblasts and DBA^+^CD45^+^ immune cells by immufluorescence. CD31^+^ endothelial cells are not DBA^+^. Their presence in our dataset might be explained by the close juxtaposition of pancreatic duct cells with endothelial cells throughout the murine pancreas (Figure S1C).

**Figure 1.**
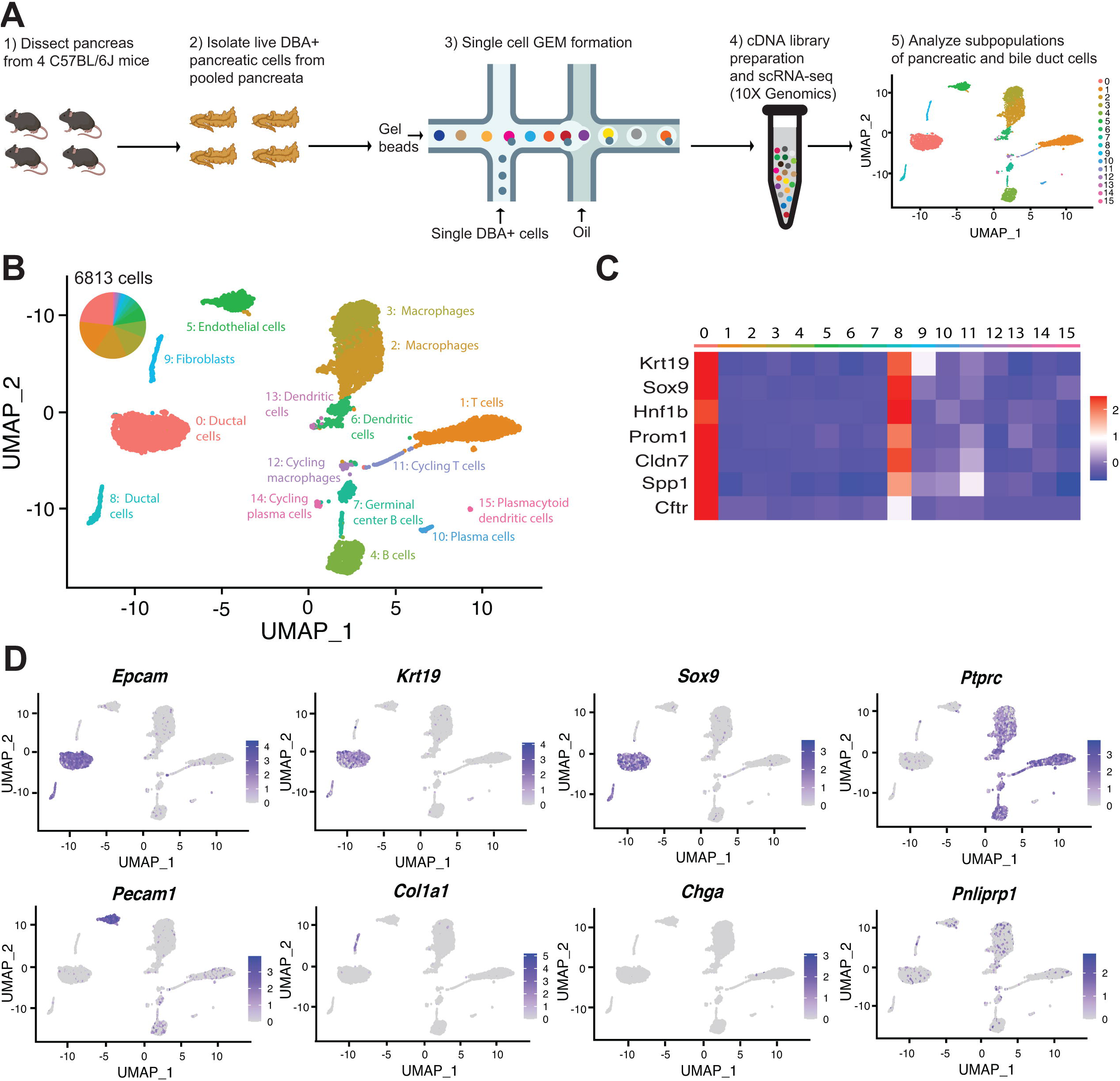
Transcriptomic map of DBA^+^ pancreatic cells. A) Schematic of experiment workflow. B) The UMAP depicts murine pancreatic DBA^+^ cells obtained using the protocol. C) A matrix plot shows average expression of ductal cell markers in all clusters, identifying clusters 0 and 8 as ductal cells. D) Feature plots illustrate markers of various cell types including epithelial (*Epcam*), ductal (*Krt19* and *Sox9*), CD45^+^ immune cells (*Ptprc*), endothelial cells (*Pecam1*), fibroblasts (*Col1a1*), endocrine cells (*Chga*), and acinar cells (*Pnliprp1*). We observed low level expression of acinar cell markers uniformly across all clusters that is likely contaminating acinar cell mRNA.

### Subpopulations of ductal cells are characterized by unique gene signatures and regulation of pathways

To get a better understanding of duct cell heterogeneity, we generated an Uniform Manifold Approximation and Projection (UMAP) plot using all duct cells (clusters 0 and 8), which revealed six distinct ductal clusters. Annotation of each duct cluster was accomplished using DEGs, Ingenuity Pathways Analysis (IPA) and upstream regulator analysis, and marker validation in murine and human pancreas (Figure 2A-D, Figure S1D-E, and Datasets S2-S4). Gene and transcript counts for each cluster are shown in Figure S1F and Dataset S1. We observed variable expression of known ductal markers within clusters. Notably, fewer murine duct cells express the transcription factor Hnf1b when compared to Sox9. This observation is in contrast to a previous report demonstrating a similar prevalence of adult murine Hnf1b^+^ and Sox9^+^ duct cells, which might be explained by different ductal cell isolation methods (Figure S1G) (13).

**Figure 2.**
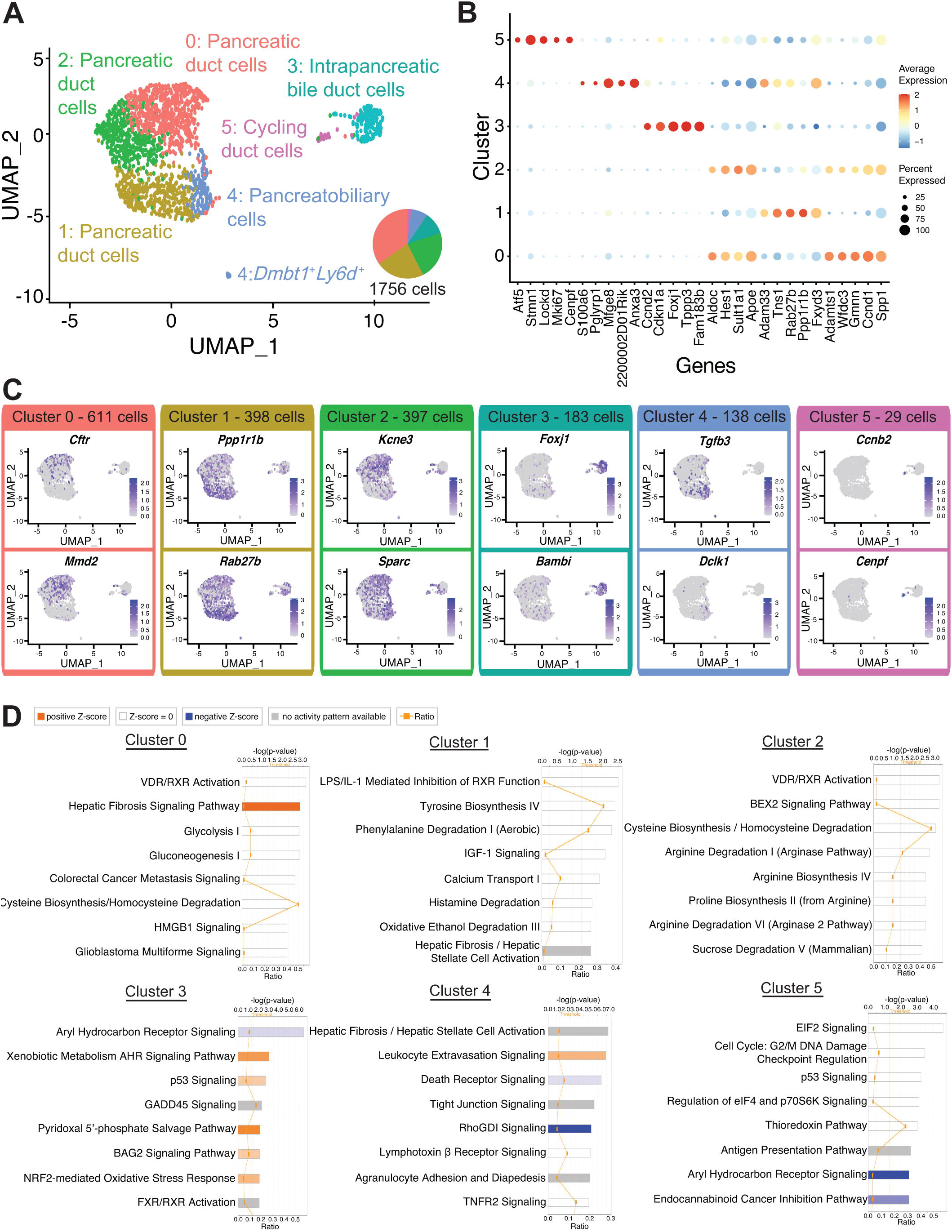
Transcriptomic map of DBA^+^ pancreatic duct cells. A) UMAP depicts identity of clusters. B) The dot plot shows the top five significantly DEGs with the highest fold change for each cluster. C) Feature plots show expression of significantly DEGs for clusters 0, 1, 3, 4, and 5. Cluster 2 is characterized by lack of or low level expression of significantly DEGs found in other clusters. D) IPA results show the top 8 deregulated pathways when comparing a cluster to all other clusters. The ratio line indicates the fraction of molecules significantly altered out of all molecules that map to the canonical pathway from within the IPA database. A positive z-score represents upregulation, and a negative z-score indicates downregulation of a pathway in that cluster when compared to all other clusters. A gray bar depicts significant overrepresentation of a pathway, the direction of which cannot yet be determined.

#### Cluster 0

Cluster 0 contains the most cells of all duct clusters in the dataset (Dataset S1). A gene that positively regulates Ras signaling *Mmd2*, the voltage-gated potassium channel protein encoded by *Kcne3*, as well as the ATP-binding cassette (ABC) transporter chloride channel protein encoded by *Cftr,* were significantly upregulated in cluster 0 when compared to all other ductal clusters (Figure 2C and Dataset S2). IPA showed upregulation of hepatic fibrosis signaling pathway in cluster 0 (Figure 2D and Dataset S3). IPA upstream regulator analysis predicted an activated state for the transcriptional regulator *Ctnnb1* in cluster 0. Predicted activation of D-glucose and growth factors Fgf2, Lep, and Hgf, suggest cluster 0 is a metabolically active duct cell subpopulation (Dataset S4). Notably, cluster 0 shows upregulation or activation of multiple genes whose alteration play important roles in the pathophysiology of pancreatic diseases such as *CFTR* for hereditary chronic pancreatitis (14) and *TGFB2* and *CTNNB1* for pancreatic cancer (15–17) (Dataset S2).

To validate gene expression patterns and determine the location of cluster 0 cells within the hierarchical pancreatic ductal tree (18), we next examined expression of select significantly DEGs. *Gmnn*, an inhibitor of DNA replication, was present in both clusters 0 and 2, so we decided to examine histologically, and were surprised to find rare protein expression of *Gmnn*, which was in contrast to the widespread RNA expression depicted by the feature plot (Figure S2A). After examining more than 1500 main pancreatic duct cells from 5 donors, we were unable to find a GEMININ positive cell, indicating very low or absent expression of *GMNN* in human main pancreatic ducts. *Spp1*, which encodes for Osteopontin, and *Wfdc3*, which are significantly DEGs in both clusters 0 and 2, show cytoplasmic expression in all mouse and human pancreatic duct types (Figure S2B-C and Table S1).

#### Cluster 1

Cells in cluster 1 have significantly upregulated expression of the exosome biogenesis gene *Rab27b* as well as *Ppp1r1b* that encodes for a molecule with kinase and phosphatase inhibition activity (Figure 2A-C and Dataset S2). IPA upstream regulator analysis predicted an activated state for the transcriptional regulator *Smarca4* and the two growth factors Tgfb1 and Gdf2 (Dataset S4). IPA results showed an enrichment in molecules regulating Calcium Transport I (Figure 2D and Dataset S3). Intracellular calcium signaling in pancreatic duct cells is an important regulator of homeostatic bicarbonate secretion (19). *PPP1R1B*, *SMARCA4*, and *TGFB1* have well described roles in the pathogenesis of pancreatic cancer (20–22). We observed expression of markers of cluster 1, *Anxa3* and *Pah*, which are also DEGs in cluster 4, to have cytoplasmic expression in all mouse and human pancreatic duct types (Figure S3A-B and Table S1). Co-staining of CFTR, a marker of cluster 0, and ANNEXIN A3 show both overlapping and non-overlapping patterns of expression in human intercalated ducts, validating the heterogeneity observed in our murine pancreatic duct dataset in human pancreatic duct cells (Figure S3C).

#### Cluster 2

Cluster 2 is characterized by low level or lack of expression of multiple ductal cell markers (*Cftr*, *Kcne3*, *Sparc*, *Mmd2, Krt7*) found in other clusters (Figure 2B-C and Figure S1G). IPA analysis showed significant overrepresentation of molecules from several pathways involved in Arginine metabolism in cluster 2 (Figure 2D and Dataset S3). Cluster 2 has the lowest average expression of total genes and transcripts (Figure S1F and Dataset 1). We therefore posit that cluster 2 represents a stable, fairly transcriptionally and metabolically inactive duct cell subpopulation when compared to other duct clusters.

#### Cluster 3

Cluster 3 cells are located almost entirely within cluster 8 of the UMAP containing 16 DBA^+^ clusters (Figure S1E). This, along with high expression of genes regulating cilia biogenesis (*Foxj1*, *Cfap44, Tuba1a*) led to the identification of cluster 3 as intrapancreatic bile duct cells (Figure 2A-C and Dataset S2). Expression of cilia biogenesis genes is more prominent in intrapancreatic bile duct cells when compared to pancreatic duct cells (Figure S3D, Dataset S2, and data not shown). IPA showed upregulation of p53 signaling, NRF2-mediated oxidative stress response, PI3K signaling in B lymphocytes, Senescence pathway among other pathways and inhibition of cell cycle: G1/S checkpoint regulation (Figure 2D and Dataset S3). IPA upstream regulator analysis predicted an activated state for transcriptional regulators *Foxa2*, *Tp53*, *Foxo3*, *Mtpn*, *Tp63*, *Smarca4*, *Nfe2l2*, *Myc*, *Tcf7l2*, *Atf4*, *Pax7*, *Smarcb1*, *Mitf*, *Sp1*, *Rel*, *Lhx1*, *Gli1*, and an inhibited state for transcriptional regulators *Mdm4*, *Gmnn*, and *Hdac1*. Five different microRNAs including mir-17 and mir-25 were all predicted to be in an inhibited state by IPA upstream regulator analysis when compared to all other duct clusters in the dataset (Dataset S4). These results highlight the differences between cellular pathways essential for homeostatic function of pancreatic duct cells and intrapancreatic bile duct cells.

#### Cluster 4

Cells in cluster 4 have significantly higher expression of *Tgfb3* and *Dclk1* when compared to all other ductal clusters (Figure 2C and Dataset S2). *Dclk1* labels tuft cells which are present in normal murine intrapancreatic bile ducts and pancreatobiliary ductal epithelium, but not in normal murine pancreatic ducts (23). *Dclk1* also marks rare normal murine pancreatic duct cells (24). Clustering analysis of *Dclk1^+^* cells in our dataset showed no subpopulations of *Dclk1^+^* ductal cells (data not shown). IPA showed 144 significantly differentially expressed pathways when comparing cluster 4 to all other ductal clusters (Figure 2D and Dataset S3). Yap, a transcriptional regulator essential for homeostasis of biliary duct cells (25), was predicted to be in an activated state by IPA upstream regulator analysis (Dataset S4). Cluster 4 also contained a small population (13 cells) of *Dmbt1* and *Ly6d*-expressing cells previously identified in extrahepatic biliary epithelium (25) (Figure S4A). These 13 cells appeared as a small population separate from other cells in cluster 4 in the UMAP (Figure 2A). Similar to the IF validation reported for extrahepatic biliary epithelial cells (BECs) (25), our IF validation of *Dmbt1* and *Ly6d*-expressing cells using Cxcl5, another marker of this subpopulation, shows a greater abundance of these cells than what would be expected given the number identified in the clustering analysis (13). It is possible that this cell type is sensitive to single cell dissociation. Cells in cluster 4 are juxtaposed to pancreatic duct cells (clusters 0, 1, and 2) in the UMAP, suggesting transcriptional commonalities with pancreatic duct cells. In addition, *Dmbt1* and *Ly6d*-expressing cells are present in cluster 4, suggesting a bile duct identity. Based on these shared features of bile and pancreas ducts, we postulate that cluster 4 contains pancreatobiliary duct cells.

#### Cluster 5

Replicating duct cells are characterized by high expression of *Mki67*, *Cenpf*, and *Cenpe* and comprise 1.65% of all duct cells in our dataset (Figure 2A-C, Figure S3D, and Dataset S2). The location of proliferating ductal epithelium is close to clusters containing intrapancreatic bile duct and pancreatobiliary cells. In addition, cluster 5 shares 6/25 DEGs with cluster 3. These data suggest that cluster 5 is comprised primarily of proliferating cluster 3 and 4 cells (Figure 2A, Figure S1E, and Dataset S2). IPA and upstream regulator analysis showed several of the same pathways and factors as those seen in clusters 3 and 4 (Datasets S3-S4). Consistent with previous reports (26, 27), pancreatic duct cells are fairly mitotically inactive.

Summarily, our high resolution single cell analysis has identified the substructure of murine pancreatic duct cells and characterized pancreatobiliary and intrapancreatic bile duct cells.

### Comparison of clusters defines heterogeneity within duct subpopulations

We next sought to determine the relationships between duct clusters by examining their similarities and differences. Dendrogram analysis, Pearson’s correlation, and DEGs revealed close relationships between clusters 0 and 2 as well as clusters 1 and 4 (Figure 3A-B and Dataset S2). Comparison of clusters 0 and 2 showed only 9 significant DEGs, suggesting a shared core gene expression program (Figure 3C-D). Overrepresentation of molecules regulating the cell cycle was observed in cluster 0 when compared to cluster 2 (Figure 3E). The DEGs upregulated in cluster 0 promote duct cell function (*Cftr*, *Tuba1a*, *Kcne3*), suggesting that cluster 0 comprises workhorse pancreatic duct cells (28).

**Figure 3.**
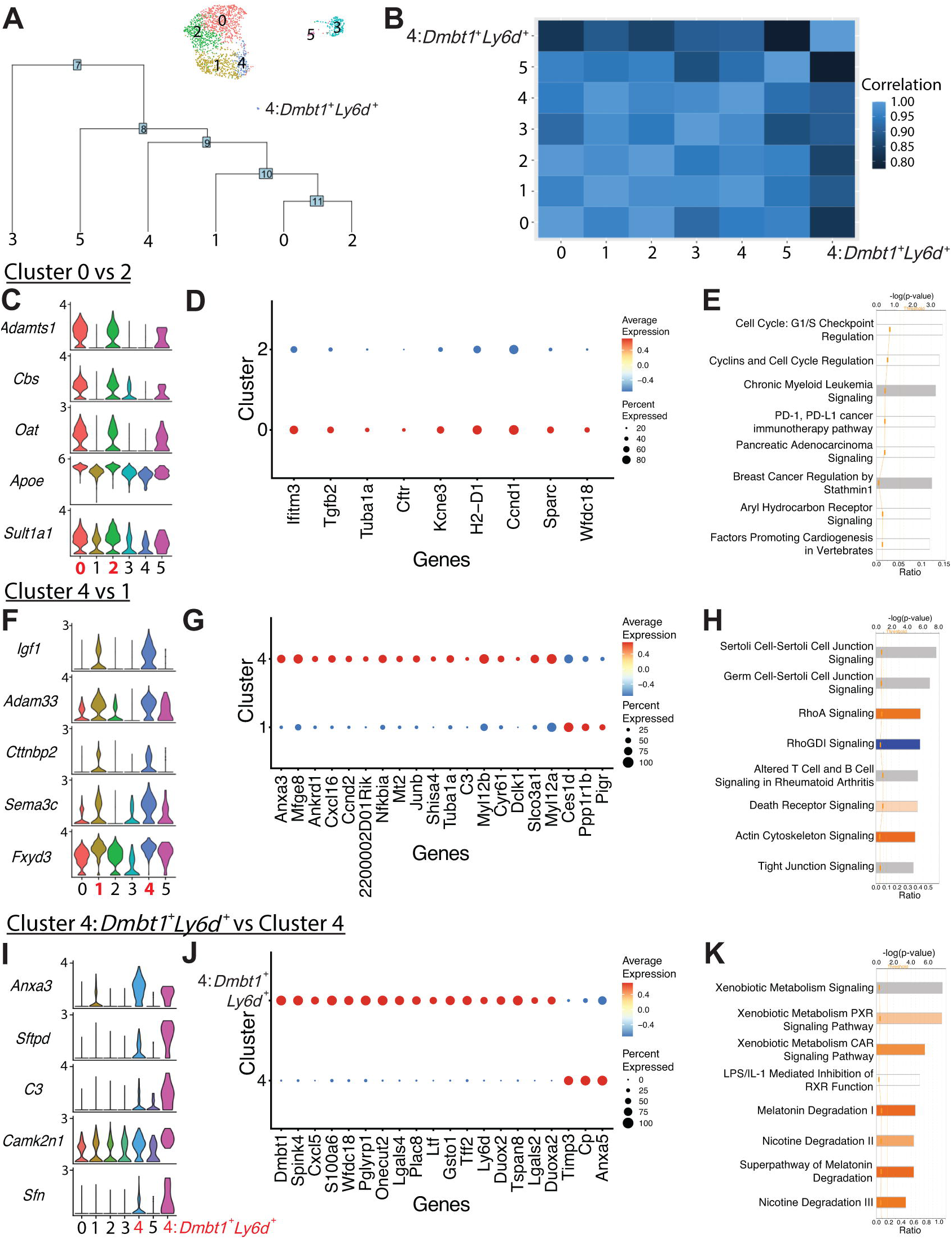
Comparison of ductal clusters 0 vs 2, 4 vs 1, and 4 vs 4: *Dmbt1^+^Ly6d^+^*. A) The cluster dendrogram created using dims (used to define the cluster) shows the Euclidean relationships between clusters. The tree is calculated in the PCA space. The genes used to define the tree were set as the variable features of the object. B) Pearson’s correlation calculated using average gene expression is depicted. C) Stacked violin plots show five DEGs sharing similar expression patterns in clusters 0 and 2. D) The dot plot shows all 9 DEGs found when comparing clusters 0 vs 2. E) The top 8 altered pathways from IPA comparing clusters 0 vs 2 are depicted. F) Stacked violin plots show five DEGs sharing similar expression patterns in clusters 4 and 1. G) The dot plot shows the top 20 DEGs ranked by fold change when comparing clusters 4 vs 1. H) The top 8 deregulated pathways from IPA comparing clusters 4 vs 1 are depicted. I) Stacked violin plots of five DEGs sharing similar expression patterns in clusters 4:*Dmbt1*^+^*Ly6d*^+^ and 4. J) The dot plot shows the top 20 DEGs ranked by fold change when comparing clusters 4:*Dmbt1*^+^*Ly6d*^+^ and 4. K) The top 8 changed pathways from IPA comparing clusters 4:*Dmbt1*^+^*Ly6d*^+^ and 4 are depicted.

When comparing pancreatobiliary cells of cluster 4 to pancreatic duct cells in cluster 1, one of the most striking differences is the enrichment in expression of genes regulating assembly of cell junctions including tight junctions, epithelial adherens junction signaling, regulation of actin-based motility by Rho, and actin cytoskeleton signaling. A strong network of stress fibers, comprised of actin filaments, myosin II, and other proteins, that function in bearing tension, supporting cellular structure, and force generation may be important for pancreatobiliary cell function and maintenance (Figure 3F-H and Datasets S3-S4) (29, 30). Cluster 4: *Dmbt1*^+^*Ly6d*^+^ cells are characterized by strong upregulation of pathways regulating Xenobiotic metabolism when compared to all other cluster 4 cells suggesting a prominent role for these cells in the bile acid and xenobiotic system (BAXS) (Figure 3I-K, and Datasets S3-S4) (31). Comparison of intrapancreatic bile duct cells and pancreatobiliary cells showed many unique features of these populations including upregulation of EIF2 signaling in pancreatobiliary cells and upregulation of coronavirus pathogenesis pathway in intrapancreatic bile duct cells (Figure S4B-D and Datasets S3-S4).

### Pancreatobiliary cells express a gene signature enriched in several targets of the Hippo signaling pathway Yap

Two subpopulations of adult murine hepatic homeostatic BECs, A and B, have been previously described (25). To determine if these subpopulations are present in intrapancreatic bile duct (cluster 3) and pancreatobiliary cells (cluster 4), we aligned our dataset with an adult hepatic murine BEC scRNA-seq dataset comprised of 2,344 homeostatic BECs (25). Intrapancreatic bile duct and pancreatobiliary cells aligned well with hepatic BECs, with no apparent batch effect (Figure S5A). Intrapancreatic bile duct cells primarily cluster together with hepatic BECs expressing subpopulation B genes, and pancreatobiliary cells primarily cluster together with hepatic BECs expressing subpopulation A genes (Figure S5B-G and Dataset S1-S2). The subpopulation A expression signature contains many genes significantly enriched as Yap targets, a signature that has been previously proposed to reflect a dynamic BEC state as opposed to defining a unique cell type (25).

### Ductal subpopulations are conserved and evident during pancreas development

To investigate whether pancreas ductal subpopulations become evident during development, we analyzed 10X Genomics single cell published datasets of epithelial-enriched pancreas cells at E12.5, E14.5, and E17.5 (32). We found distinct subpopulations of ductal cells that notably overlap in expression of key marker genes associated with adult pancreas ductal subpopulations (Figure S6A-L). As we expected, clear patterns of marker gene expression associated with adult clusters manifest at later stages of development (Figure S6D,H,L). Since the developmental biology samples were obtained from Swiss Webster mice, our results suggest the subpopulations of adult pancreas duct cells we describe in C57BL/6J mice are conserved.

### DBA^+^ lectin murine pancreas sorting identifies previously missed ductal subpopulations

To determine the novelty of adult duct cell heterogeneity manifested using DBA^+^ lectin sorting of murine pancreas, we next compared our DBA^+^ murine pancreatic ductal clusters to previously reported subpopulations of mouse and human pancreas duct cells. Using inDrop and an islet isolation pancreas preparation, Baron *et al.* (2016) identified the substructure of mouse and human pancreatic duct cells (9). Two subpopulations of mouse pancreatic duct cells characterized by expression of *Muc1* and *Tff2* (subpopulation 1) and *Cftr* and *Plat* (subpopulation 2) were described. While *Cftr* expression is characteristic of our cluster 0 (Figure 2C), *Muc1*, *Tff2*, and *Plat* expression didn’t typify any murine DBA^+^ pancreatic duct subpopulation (Figure S6M). Two subpopulations were similarly described for human pancreas duct cells characterized by expression of 1) *TFF1*, *TFF2*, *MUC1, MUC20,* and *PLAT* and 2) *CFTR* and *CD44*. *Tff1* is not expressed in murine DBA^+^ ductal cells (clusters 0-5). *Cd44* is significantly upregulated in pancreatobiliary cells, and *Muc20* as well as *Tff2* are significantly upregulated in 4:*Dmbt1*^+^*Lyd6*^+^ cells (Dataset S2 and Figure S6M-N). Dominic Grün *et al*. (2016) previously reported 4 subpopulations of human pancreatic duct cells characterized by expression of *CEACAM6*, *FTH1*, *KRT19*, and *SPP1* using an islet isolation pancreas preparation and the CEL-seq protocol (10). While *Spp1* is significantly upregulated in DBA^+^ pancreas duct clusters 0 and 2, *Fth1* doesn’t characterize any murine DBA^+^ pancreas duct population, and *Krt19* is significantly upregulated in pancreatobiliary cells (Dataset S2, Figure S1G, and Figure S6O). *CEACAM6* has no mouse homolog. The differences in pancreatic ductal subpopulation identification may be due to single cell methodology (inDrop, CEL-seq, and 10X Genomics), pancreas preparation method (islet isolation vs DBA^+^ lectin sorting), differences in ductal cell numbers analyzed, or potential differences between mouse and human duct cells.

Six subpopulations of human pancreatic duct cells have been described using the 10X Genomics platform based on sorting for *BMPR1A*/*ALK3* (8). Using AddModuleScore in Seurat, we calculated a score comparing each of our murine duct clusters to the human ALK3^+^ clusters (Figure S7A-F) (33). Murine pancreatic duct clusters 0-2 had the highest scores when compared to human ALK3^+^ clusters 1 (*SPP1*^+^ Stress/harboring progenitor-like cells) and 2 (*TFF1*^+^ activated/migrating progenitor cells). Murine pancreatobiliary cells (cluster 4) scored the highest when compared to the human ALK3^+^ cluster 3 (AKAP12^+^ small ducts). The human ALK3^+^ cluster 4 (*WSB1*^+^ centroacinar cells) didn’t distinguishably overlap with any DBA^+^ mouse pancreas ductal clusters. DBA is expressed in murine centroacinar/terminal ducts as early as three weeks of age (34), thus these cells would be expected to be present in our dataset(11). Examination of centroacinar/terminal ductal cell markers *Hes1* (35), *Aldh1a1* (36), and *Aldh1b1* (37) showed broad expression enriched in either clusters 0 and 2 (*Hes1* and *Aldh1b1*) or clusters 1 and 4 (*Aldh1a1*), rather than a distinct subpopulation as is seen in the ALK3^+^ human pancreatic duct dataset. *Aldh1a7* is negligibly expressed in murine duct clusters 0-5 (Figure S7G). Unlike in mouse DBA^+^ pancreas ductal clusters, the human ALK3^+^ dataset contains two ducto-acinar subpopulations characterized by expression of genes enriched in acinar cells. To assess the presence of ducto-acinar cells in adult murine pancreas, we performed immunolabeling for markers of the ALK3^+^ human ducto-acinar clusters 5 (*CPA1*) and 6 (*AMY2A* and *AMY2B*). Although ducto-acinar cells, like centroacinar/terminal ductal cells, don’t define a unique cluster in our DBA^+^ murine duct subpopulations, we identified DBA^+^Cpa1^+^ and DBA^+^α-amylase^+^ ducto-acinar cells in adult murine pancreas (Figure S7H). Taken together, these data suggest murine centroacinar/terminal ductal and ducto-acinar cells are largely transcriptionally homogenous with other murine duct cell types.

### RaceID3/StemID2 suggest murine DBA^+^ duct cluster 0 and 2 cells are the most progenitor-like

Given the close relationships observed between DBA^+^ duct clusters 0 and 2 as well as 1 and 4, we next assessed differentiation potential using RaceID3/StemID2 to predict cell types, lineage trajectories, and stemness (38). Unsupervised clustering with RaceID3 showed 17 clusters. RaceID3 clusters with 10 cells or less were removed from subsequent analyses, and Seurat duct clusters 3 and 5 are not included in this analysis (Figure 4A-B). RaceID3 clusters with the highest StemID2 score correlate to cells present in Seurat duct clusters 0 and 2 (Figure 4C and Figure S8A-B). The variable StemID2 scores observed for cells within Seurat duct clusters 0, 1, 2, and 4 suggest distinct stages of differentiation or maturation. Consistent with previous literature, the pancreatic ductal cell progenitor niche isn’t restricted to a single cluster (8).

**Figure 4.**
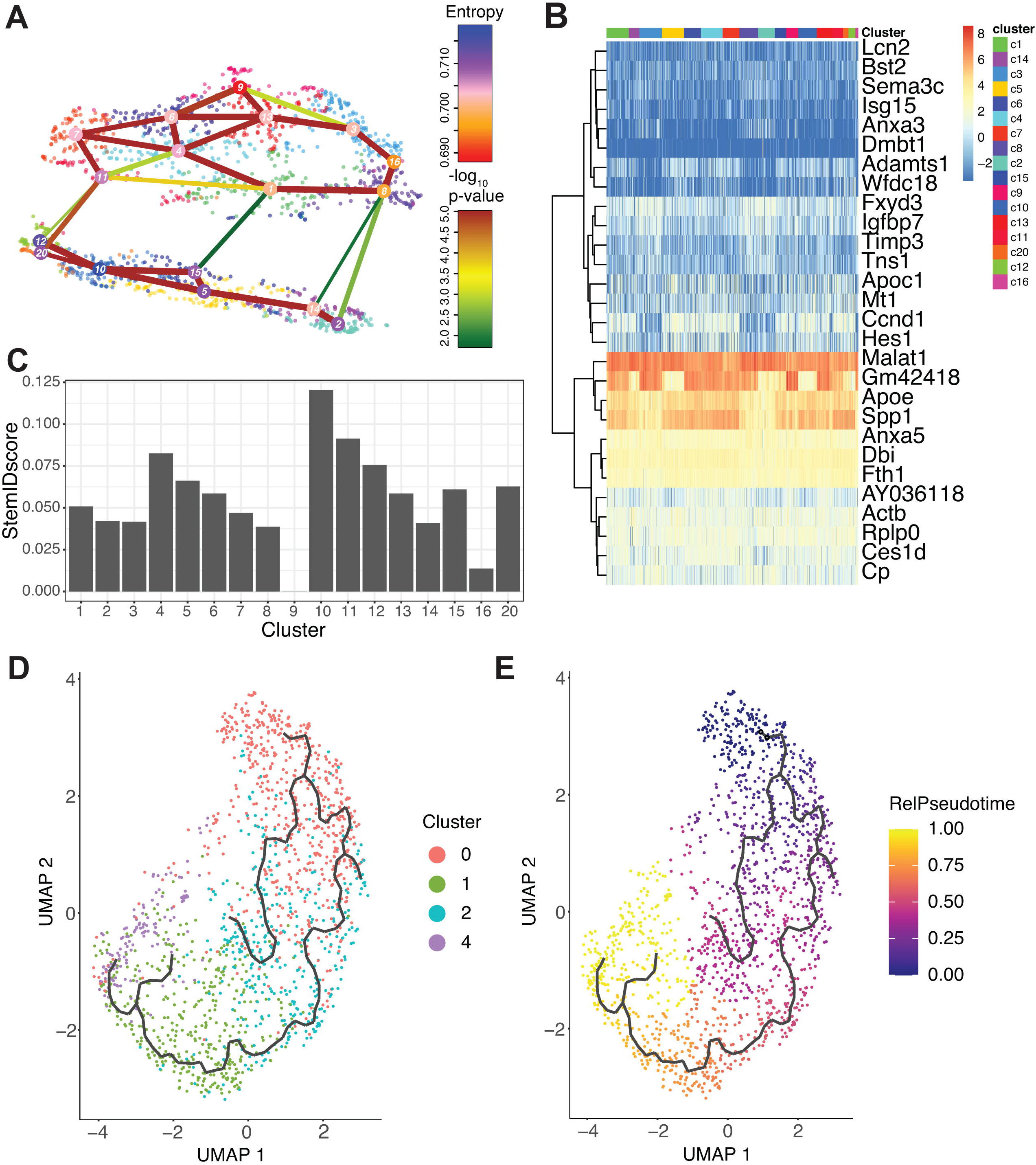
RaceID3/StemID2 predict clusters 0 and 2 have the highest progenitor potential. A) The lineage tree inferred by StemID2 is shown in the RaceID3 clusters. Node color represents the level of transcriptome entropy, edge color describes level of significance, and edge width describes link score. B) Heat map depicts expression of top 5 DEGs in RaceID3 clusters with FDR < 0.01 and fc > 1.2. C) StemID2 scores for RaceID3 clusters are graphed. D) Monocle 3 clustering of murine DBA^+^ duct clusters 0, 1, 2, and 4 are depicted. E) Each cell’s relative pseudotime value is depicted that is a measurement of the distance between its position along the trajectory and the starting point (cluster 0).

### Pseudotime ordering identifies an epithelial-mesenchymal transition (EMT) axis in pancreatic duct cells

To further examine the lineage relationships among pancreas duct subpopulations, we ordered cells in pseudotime based on their transcriptional similarity (39). Monocle 3 analysis showed DBA^+^ duct clusters 3 and 5 were disconnected from the main pseudotime trajectory, so we focused our analysis on DBA^+^ duct clusters 0, 1, 2, and 4 (Figure S8C). Because RaceID3/StemID2 analysis showed Seurat clusters 0 and 2 have the highest StemID scores, we started the pseudotime ordering beginning with cluster 0 as Seurat clusters 0 and 2 are juxtaposed in the Monocle 3 clustering (Figure 4D-E and Figure S8D).

In Monocle 3 analysis, genes with similar patterns of expression that vary over time across the pseudotime trajectory are coalesced into modules (Figure 5A). We performed IPA and upstream regulator analysis, a pairwise comparison, comparing select clusters within a module to analyze the gene expression changes along the pseudotime trajectory (Figure 5B-D and Datasets S3-S5). Examination of pathways deregulated in modules 4 and 14 showed a shift in the molecules driving the Xenobiotic Metabolism CAR Signaling Pathway. The Xenobiotic nuclear receptor CAR is an important sensor of physiologic toxins and plays a role in their removal (40). This pathway is regulated by *Aldh1b1*, *Aldh1l1*, *Gstt2*/*Gstt2b*, *Hs6st2*, and *Ugt2b7* in clusters 0 and 2 and *Aldh1a1*, *Fmo3*, *Gstm1*, and *Sod3* in cluster 1, suggesting that these clusters might respond differently when exposed to toxins or play heterogenous roles in endogenous toxin elimination (Figure 5B-C).

**Figure 5.**
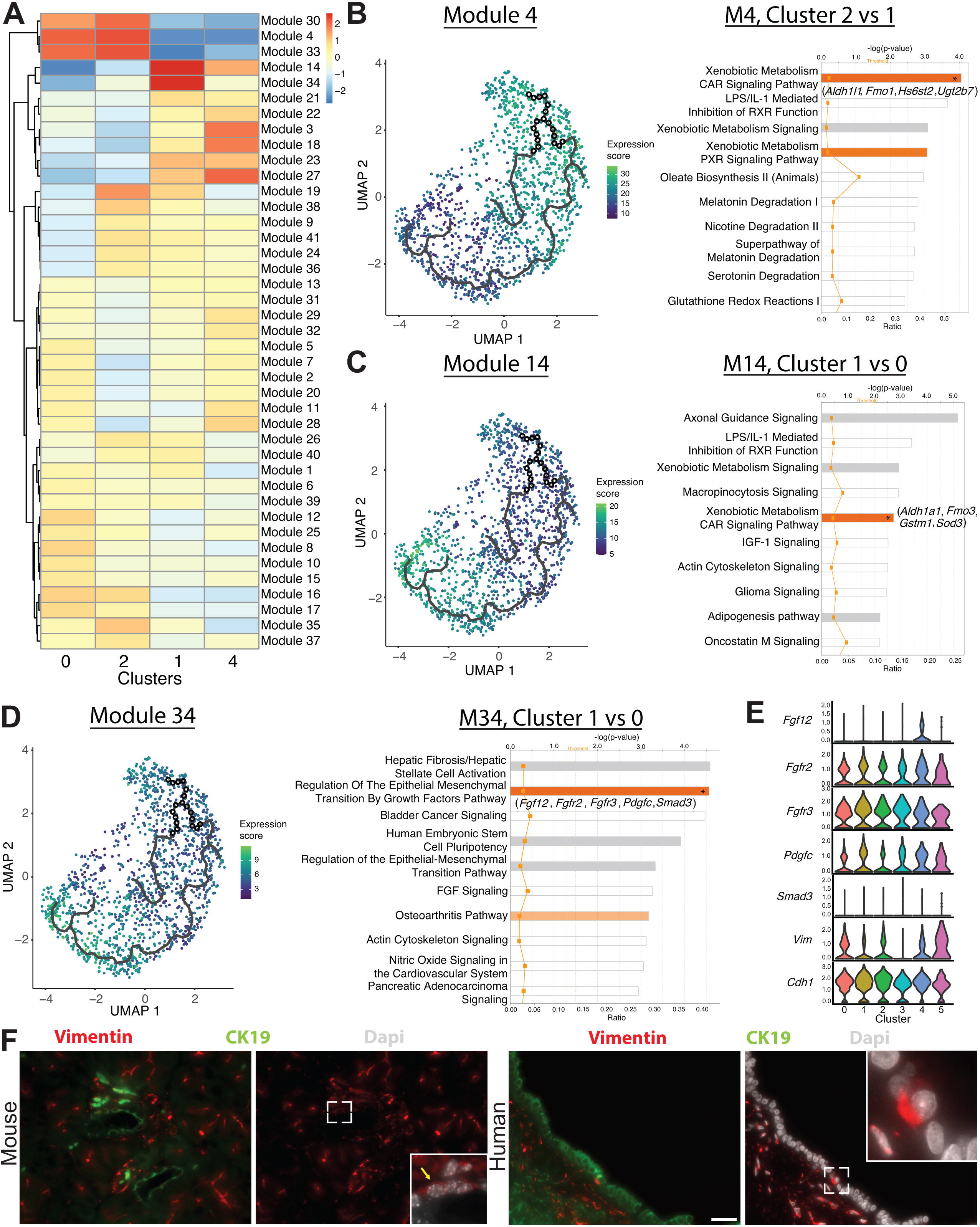
Monocle 3 analysis reveals an epithelial mesenchymal axis in pancreatic duct cells. A) Expression changes of the modules generated by Monocle 3 analysis are shown for each cluster. B-D) Expression of modules 4, 14, and 34 along with select IPA results of the top 10 deregulated pathways are shown. Genes in parenthesis are altered in the pathway containing an asterisk in the bar. E) Stacked violin plots show expression of genes in the Regulation of the Epithelial Mesenchymal Transition By Growth Factors Pathway in DBA^+^ duct clusters 0-5. F) IF depicts CK19^+^ Vimentin^+^ copositive pancreatic duct cells in mouse (yellow arrow) and human. The main pancreatic duct is shown for humans. Scale bars are 50uM.

Regulation of the Epithelial Mesenchymal Transition By Growth Factors Pathway was upregulated in cluster 1 when compared to cluster 0 in Module 34. Molecules altered in this pathway play variable roles in promoting the epithelial or mesenchymal state and include *Fgf12*, *Fgfr2*, *Fgfr3*, *Pdgfc*, and *Smad3* (Figure 5D). When comparing clusters 0 and 1, examination of EMT markers *Vim* and *Cdh1* showed a stronger probability of expression of *Cdh1* in cluster 1 and a stronger probability of expression of *Vim* in cluster 0 (Figure 5E). Using immunofluorescence (IF), we detected Vimentin^+^ ductal cells in both mouse and human pancreas, validating this epithelial-mesenchymal transitional axis (Figure 5F).

### *SPP1* is required for mature human pancreas duct cell identity

Our analysis thus far has identified and validated multiple transcriptional programs expressed by murine pancreatic duct cells and predicted possible lineage relationships among them. To assess the function of select markers defining murine DBA^+^ pancreas duct clusters, we next examined the consequences of their loss in the immortalized human duct cell line HPDE E6/E7 (hereafter HPDE). *Spp1*, a marker of clusters 0 and 2, has been shown by us and others to mark a pancreas duct cell type enriched in progenitor capacity (8, 41). *Gmnn*, a marker of cluster 0, acts to inhibit re-replication of DNA during DNA synthesis by inhibiting the prereplication complex (42, 43). *Anxa3*, a marker of clusters 1 and 4, inhibits phospholipase A2 and cleaves inositol 1,2-cyclic phosphate generating inositol 1-phosphate in a calcium dependent manner (44, 45). *PAH* and *WFDC3* were not expressed in HPDE cells (data not shown). We generated and validated *SPP1*, *GMNN*, and *ANXA3* knockout HPDE lines using CRISPR/Cas9 (Figure 6A-C). Strong, consistent phenotypes were observed among different knockout lines for each gene despite some lines not demonstrating full loss of the protein (HPDE *ANXA3* gRNA2 and HPDE *SPP1* gRNAs 1-4). Cellular morphology was similar to the scrambled (scr) gRNA control (46) for every knockout line except HPDE *SPP1* gRNAs 1-4, which displayed a dramatic change in cellular morphology. HPDE *SPP1* knockout cells showed prominent filipodia and significantly increased proliferation in both 2D and 3D assays. HPDE *ANXA3* and *GMNN* knockout lines also show significantly increased proliferative capacity in both 2D and 3D assays, a phenotype suggestive of increased progenitor function (Figure 6D-F, Figure S9A, and data not shown). The change in cellular morphology in HPDE *SPP1* knockout lines is accompanied by decreased duct function as measured by carbonic anhydrase activity (Figure 6G).

**Figure 6.**
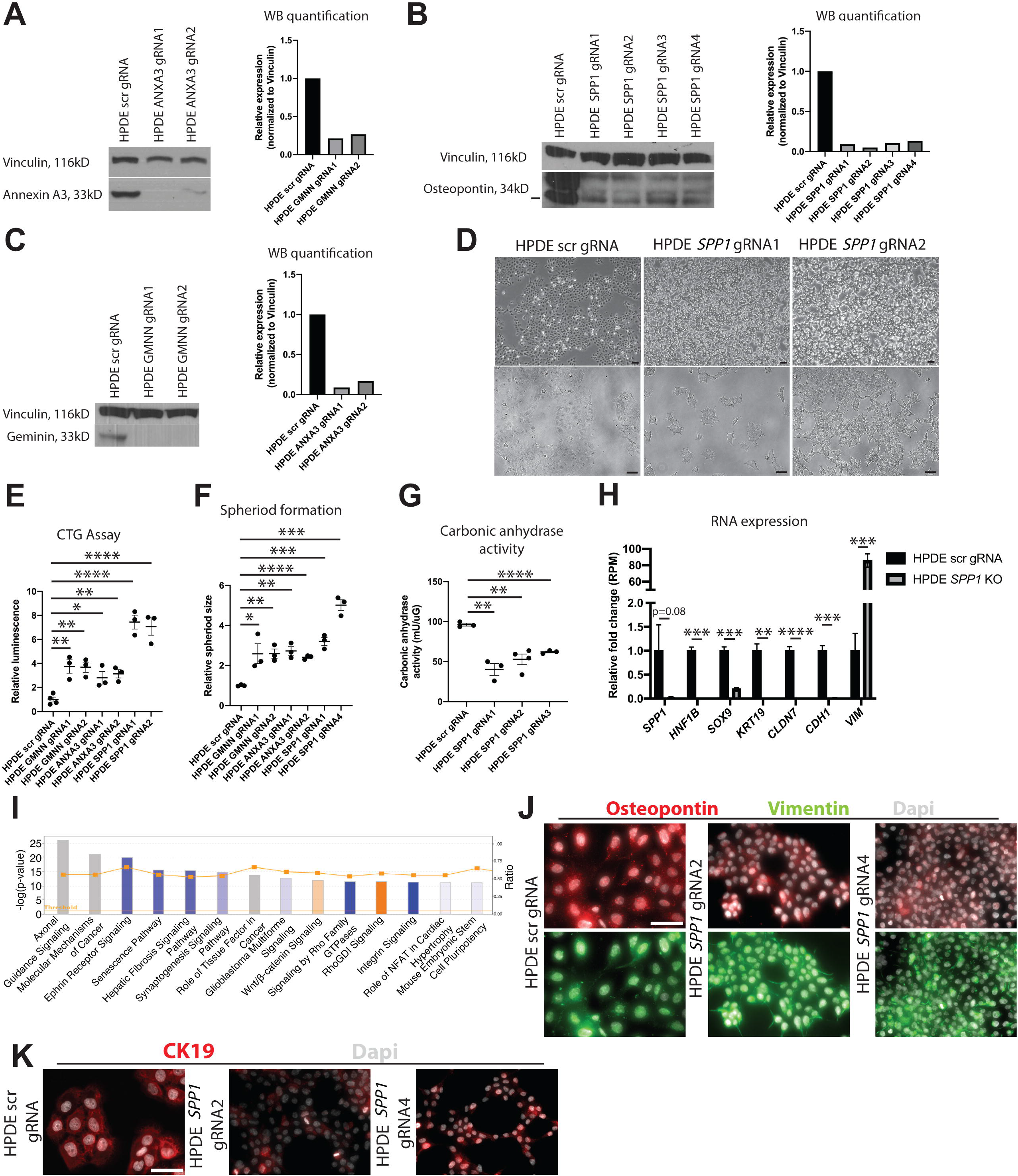
SPP1 loss promotes a progenitor-like state in human pancreatic duct cells. A-C) Western blot and quantification of western blot images shows expression of ANNEXIN A3, OSTEOPONTIN, and GEMININ in knockout HPDE E6/E7 lines and the control. D) Brightfield images show changes in cellular morphology of HPDE E6/E7 *SPP1* knockout lines. Scale bars are 100 μm. E) Proliferation was assessed in 2D by CTG assays. F) Proliferation was assessed in 3D by seeding 1000 cells in a drop of matrigel. G) Significantly decreased carbonic anhydrase activity is observed in HPDE E6/E7 *SPP1* knockout lines when compared to the control. H) Relative fold changes calculated using RPM values of mesenchymal and duct markers are shown. Average RPM values for *SPP1* are 79.8 ± 43.2 (scr) and 2.8 ± 0.4 (KO), *HNF1B* are 418.5 ± 33.4 (scr) and 1 ± 0.3 (KO), *SOX9* are 1,555.2 ± 124.8 (scr) and 316 ± 38.7 (KO), *KRT19* are 16,789.5 ± 2,431.2 (scr) and 61 ± 60.5 (KO), *CLDN7* are 8,651.8 ± 923.2 (scr) and 112.8 ± 27.7 (KO), *CDH1* are 8,651.8 ± 923.2 (scr) and 112.8 ± 27.8 (KO), and *VIM* are 80.2 ± 29.1 (scr) and 6,879.8 ± 652.6 (KO). I) The top 14 deregulated pathways from IPA are shown comparing HPDE E6/E7 *SPP1* KO vs HPDE E6/E7 scr gRNA control. J) Immunocytochemistry (ICC) demonstrated reduced Osteopontin expression in HPDE E6/E7 *SPP1* gRNA2 and HPDE E6/E7 *SPP1* gRNA4 when compared to HPDE E6/E7 scr gRNA. Vimentin ICC depicts organized intermediate filaments in HPDE E6/E7 *SPP1* gRNA2 and HPDE E6/E7 *SPP1* gRNA4 while HPDE E6/E7 scr gRNA cells show diffuse, light labeling. Scale bar is 50 μm. K) CK19 ICC shows organized intermediate filaments in HPDE E6/E7 scr gRNA cells while HPDE E6/E7 *SPP1* gRNA2 and HPDE E6/E7 *SPP1* gRNA4 cells display punctate CK19 labeling, where present. Scale bar denotes 50 μm.

To assess the changes in HPDE *SPP1* knockout lines on a molecular scale, we performed bulk RNA-sequencing on all 4 HPDE *SPP1* knockout lines and the HPDE scr gRNA control. A significant increase in markers associated with epithelial-mesenchymal transition (EMT) (*VIM*, *ZEB1*, *TWIST1*, *MMP2*) was observed in HPDE *SPP1* knockout lines when compared to the control (Datasets S2-S4 and Figure 6H-I). Markers of mature duct cells (*HNF1B*, *SOX9*, *KRT19*) were significantly downregulated in HPDE *SPP1* knockout lines when compared to the control (Figure 6H, J-K and Dataset S2). Gene Set Enrichment Analysis (GSEA) showed positive enrichment of pathways that regulate embryogenesis (*HOX* genes and *NOTCH* signaling) and cell cycle regulation in HPDE *SPP1* knockout lines when compared to the HPDE scr gRNA control, supporting the notion that loss of SPP1 leads to a more immature, progenitor-like state (Figure S9B-E). Taken together, these results define unique functional properties for markers that characterize murine DBA^+^ pancreas duct cells and suggest that *SPP1* is an essential regulator of human pancreatic duct cell maturation and function.

Transdifferentiation of pancreatic duct cells to endocrine cells at early postnatal stages and in pancreatic injury models has been suggested by several studies (47, 48). To query whether HPDE SPP1 knockout progenitor-like, dedifferentiated duct cells harbor the capacity to re-differentiate to endocrine cells *in vivo*, we injected HPDE SPP1 knockout cell lines and HPDE scr gRNA control cells subcutaneously into NSG mice (Figure 7A). After 5 days post-injection, α-amylase^+^ CK19^+^ double positive cells were evident in HPDE scr gRNA control cells, but not in HPDE SPP1 knockout cells (Figure 7B). This observation is consistent with the previously described ducto-acinar axis characteristic of human pancreatic duct cells (8). We observed Ngn3^+^Sox9^+^ copositive HPDE SPP1 knockout cells, suggesting potential for differentiation towards the endocrine lineage (Figure 7C). We detected Synaptophysin^+^ Glucagon^+^ as well as Synaptophysin^+^ C-peptide^+^ double positive HPDE SPP1 knockout cells (Figure 7D-E). Expression of endocrine markers in subcutaneously injected HPDE scr gRNA cells was not observed at Day 5 post-injection. A small subset of C-peptide^+^ HPDE SPP1 knockout cells express both Pdx1 and Nkx6.1 (Figure 7E-F). Together, these data point to previously unappreciated roles for SPP1 in maintaining duct cell properties and preventing changes in cell identity.

**Figure 7.**
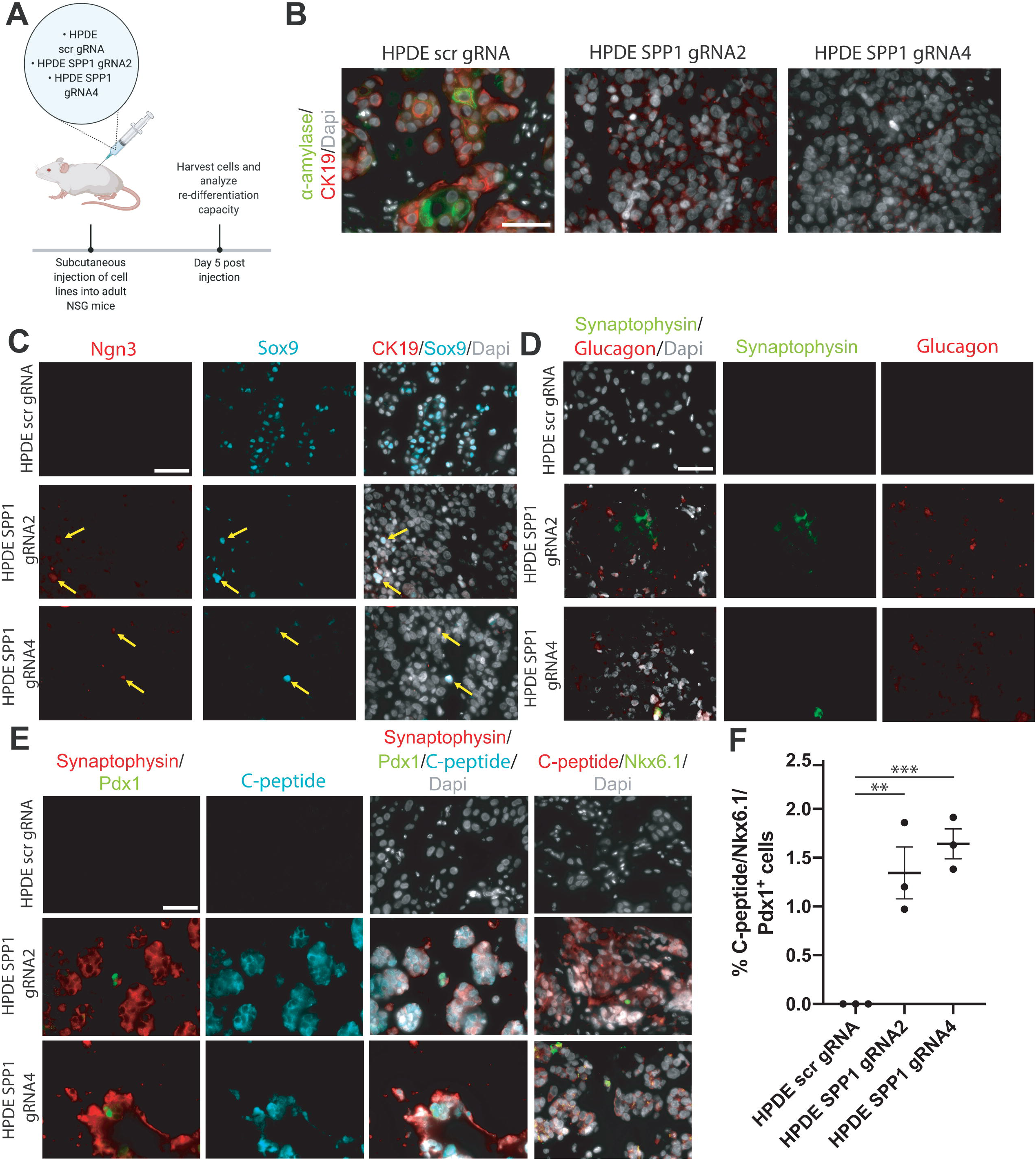
HPDE SPP1 knockout cells are capable of differentiating into cells with endocrine appearance, including cells exhibiting α- and β-like appearance, but not duct-like or acinar-like cells *in vivo.* A) Schematic of *in vivo* experiment. B) IF shows CK19^+^ α-amylase^+^ double positive HPDE scr gRNA cells. C) IF depicts Ngn3^+^ Sox9^+^ double positive HPDE SPP1 knockout cells (yellow arrows). D) Synaptophysin^+^ Glucagon^+^ double positive cells (yellow arrows) are detected in HPDE SPP1 knockout cells. E) C-peptide, Nkx6.1, and Pdx1 expression are evident in HPDE SPP1 knockout cells. F) The percentage of C-peptide^+^, Nkx6.1^+^, and Pdx1^+^ triple positive cells for HPDE scr gRNA cells is 0, HPDE SPP1 gRNA2 is 1.343 ± 0.2664, and HPDE SPP1 gRNA 4 cells is 1.642 ± 0.1533. *All* scale bars in this figure are 50 μm.

### Geminin safeguards against accumulation of DNA damage in mouse ductal cells in the setting of chronic pancreatitis

One marker of the workhorse population of pancreatic duct cells *Gmnn* has previously been associated with chronic inflammatory diseases such as asthma (49). We therefore queried its role in pancreas inflammatory disease. *Gmnn* binds to Cdt1 and inhibits DNA replication during the S phase. Geminin is a crucial regulator of genomic stability; its inhibition in multiple cancer cell lines leads to DNA re-replication and aneuploidy (50, 51). To determine the requirement for *Gmnn* in normal homeostatic pancreatic ductal cells, we generated a conditional *Gmnn* floxed allele and crossed the mouse to the *Sox9–Cre^ERT^*^2^ (52) and *Hnf1b–Cre^ERT^*^2^ (53) lines (Figure S10). Adult mice, between the ages of 7-9 weeks, were injected with tamoxifen to ablate Geminin in mouse pancreatic duct cells. Tamoxifen injected *Sox9cre^Tg/wt^*; *Geminin^f/f^*, *Sox9cre^Tg/wt^*; *Geminin^f/wt^*, and *Hnf1b^Tg/wt^*; *Geminin^f/f^* mice displayed no histological abnormalities as assessed by hematoxylin and eosin (H&E) staining and no significant alterations in DNA damage as assessed by ATR and γ-H2AX IF up to 6 months post tamoxifen injection (data not shown). We were unsurprised by these findings, given the low proliferation rate of murine pancreatic duct cells suggested by our single cell data. Thus, Geminin may only be required in the context of pathologies characterized by increased proliferation in the pancreas such as pancreatitis or PDA (54).

We examined proliferation in human pancreas duct cells in CP patients (N=5 patients) and found a significant increase in Geminin expression when compared to normal human pancreatic duct cells (N=10 donors) (Figure 8A-B). Pancreatic duct ligation (PDL), an experimental technique that recapitulates features of human gallstone pancreatitis, results in an increase in proliferation of rat pancreatic duct cells (55, 56). To investigate the role of Geminin in mouse pancreatic duct cells in the setting of CP, we performed PDL on *Sox9cre^Tg/wt^*; *Geminin^f/f^*, *Sox9cre^Tg/wt^*; *Geminin^f/wt^*, *Hnf1b^Tg/wt^*; *Geminin^f/f^* and littermate control mice (Figure 8C). As in the human setting, we also observed upregulation of Geminin in ductal epithelium in the control PDL mouse group (Figure 8D). Previously reported features of the PDL model were evident in our transgenic mice including replacement of parenchymal cells with adipose tissue, inflammation, and fibrosis (57, 58) (Figure S11A-B). Significant attenuation of Geminin expression was observed in *Sox9cre^Tg/wt^*; *Geminin^f/f^*, *Sox9cre^Tg/wt^*; *Geminin^f/wt^*, and *Hnf1b^Tg/wt^*; *Geminin^f/f^* mouse pancreatic duct cells when compared to controls (Figure 8D and Figure S12A). Homozygous *Gmnn* loss in Sox9^+^ pancreatic ductal cells promoted an acute increase in proliferation, as assessed by BrdU incorporation, at Day 7 which became insignificant at Day 30 (Figures S12B-E). No changes were observed in apoptosis for any model or time point when compared to controls as assessed by cleaved caspase-3 IF (data not shown). Examination of DNA damage by γ-H2AX IF showed significantly increased γ-H2AX foci in *Sox9cre^Tg/wt^*; *Geminin^f/f^* mice at Day 7, an observation that was sustained at Day 30 (Figure 8E-H). Assessment of DNA damage in *Sox9cre^Tg/wt^*; *Geminin^f/f^*, *Sox9cre^Tg/wt^*; *Geminin^f/wt^*, and *Hnf1b^Tg/wt^*; *Geminin^f/f^* mice by ATR IF showed no significant changes (data not shown). The lack of phenotypes observed in the *Hnf1b^Tg/wt^*; *Geminin^f/f^* model may be due to differences in recombination induced by the *Sox9–Cre^ERT^*^2^ and *Hnf1b–Cre^ERT^*^2^ lines, since fewer cells of the pancreatic ductal epithelium express Hnf1b (Figure S1G and Figure 8C). Taken together, these data suggest Geminin is an important regulator of genomic stability in pancreatic ductal cells in the setting of CP.

**Figure 8.**
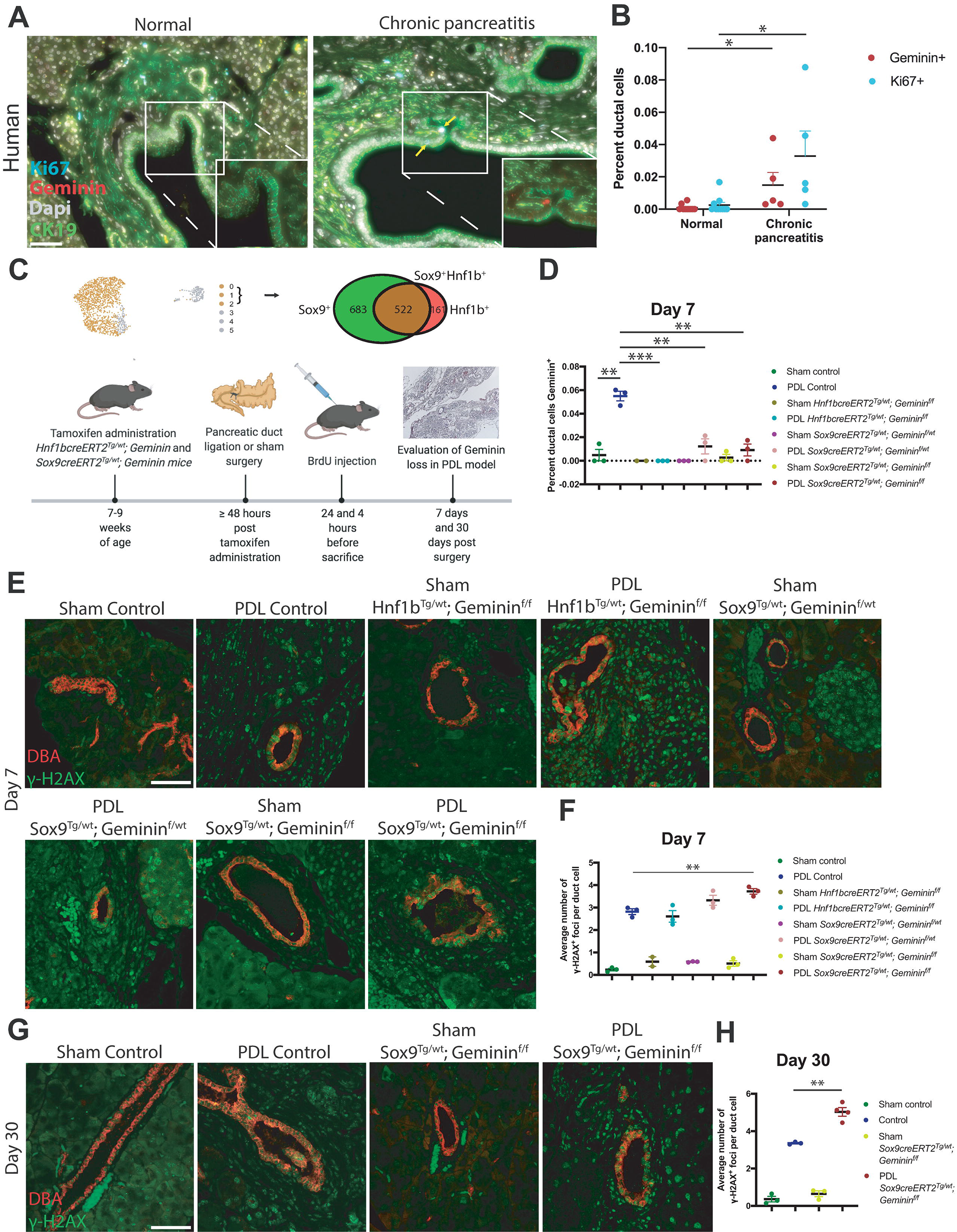
*Gmnn* is a regulator of genomic stability in mouse pancreatic duct cells during chronic pancreatitis. A-B) IF images and quantification show a significant increase in proliferation in pancreatic duct cells in CP patients when compared to normal human pancreatic duct cells. C) A schematic of tamoxifen and BrdU administration is shown. The UMAP depicts the pancreatic cells (clusters 0-2) that were analyzed in this experiment. The Venn diagram shows the number of cells in clusters 0-2 that are Sox9^+^, Hnf1b^+^, and Sox9^+^Hnf1b^+^ copositive. D) Quantification of Geminin positive ductal cells at Day 7 in *Sox9cre^Tg/wt^*; *Geminin^f/f^*, *Sox9cre^Tg/wt^*; *Geminin^f/wt^*, *Hnf1b^Tg/wt^*; *Geminin^f/f^* and control mice is depicted. E-F) Representative IF images and quantification of γ-*H2AX positive foci is shown at Day 7 in the PDL transgenic models. G-H)* Representative IF images and quantification of γ-*H2AX positive foci are shown at Day 30 in the Sox9cre^Tg/wt^*; *Geminin^f/f^* and control PDL models*. All* scale bars in this figure are 50 μm.

## DISCUSSION

We present a single cell transcriptional blueprint of murine pancreatic duct cells, intrapancreatic bile duct cells, and pancreatobiliary cells. Notably, our single cell analysis showed that endothelial cells, fibroblasts, and immune cells are also obtained using the DBA^+^ lectin sorting strategy(12), and suggests that a subsequent ductal purification step is required to obtain pure pancreatic duct cells using this protocol. A static transcriptional picture in time has highlighted a very dynamic view of pancreas duct cell heterogeneity. Our study provokes reinterpretation of several previously published lineage tracing reports using ductal-specific Cre mouse lines, and will help plan future lineage tracing studies.

Cluster 0 workhorse pancreatic duct cells comprise the largest pancreatic duct subpopulation identified. Although clusters 0 and 2 share many markers, we found compelling differences in metabolic states as manifested in part by an overall lower gene and transcript count for cluster 2. IPA suggested that subpopulations of pancreatic duct cells may use different predominant mechanisms for bicarbonate secretion such as Cftr (59) for cluster 0 and calcium signaling for cluster 1 (60). One notable difference between clusters 0 and 2 vs 1 is the molecules which regulate the Xenobiotic Metabolism CAR signaling pathway. We observed expression of several genes, whose alteration contributes to PDA progression including *Tgfb2* and *Ctnnb1* in cluster 0 and *Ppp1r1b, Smarca4,* and *Tgfb1* in cluster 1. IPA upstream regulator analysis of Monocle 3 Module 14 predicted significant inhibition of *Kras* in cluster 1 when compared to cluster 0. Additionally, IPA upstream regulator analysis comparing cluster 2 vs 0 in Module 19 predicted activation of Myc and Mycn in cluster 2. These genes play central roles in homeostasis of pancreatic duct cells, and it’s possible that distinct ductal cell subpopulations which are actively expressing these pathways may have different predispositions to PDA with mutations in these genes, heterogeneity which may also contribute to development of different subtypes of PDA.

The role of SPP1 in homeostatic pancreatic ductal cells has been elusive, since *Spp1* knockout mice have no apparent pancreatic duct phenotypes (41). We identified an EMT axis in pancreatic duct cells using Monocle 3 and validated this observation in mouse and human duct cells. *Spp1* is one gatekeeper of this epithelial to mesenchymal transitory duct phenotype as manifested by loss of ductal markers, reduced duct function, and upregulation of EMT genes in HPDE *SPP1* knockout cells when compared to controls. Clusters 0 and 2, characterized by strong expression of *Spp1*, show the highest StemID2 scores. *SPP1* knockout HPDE cells display prominent filipodia and the highest proliferative capacity of all markers examined when compared to controls. Taken together, these phenotypes along with upregulation of pathways regulating mammalian development (Notch signaling and Hox genes) manifested by GSEA suggest SPP1 loss promotes human duct cell de-differentiation.

During pancreas development, the multipotent epithelial progenitors become increasingly compartmentalized into tip and trunk progenitors that give rise to acinar and endocrine/ductal cells, respectively (61). Our data suggest that SPP1-deficient HPDE cells dedifferentiate into a trunk, and not tip, progenitor-like cell and that re-differentiation of HPDE cells to a human pancreatic duct or acinar cell lineage isn’t favored *in vivo* following SPP1 loss. These data underscore the requirement for SPP1 expression for mature human pancreas duct cell identity. It has been hypothesized that SPP1’s role in mature pancreatic duct cells is evident during pathogenesis. Several groups have already nicely shown that Spp1 plays important roles in pancreatic pathologies including PDAC (62, 63). In human pancreas duct cells, the subpopulation characterized by *SPP1* expression is described as “stress/harboring progenitor-like cells” (8). We observed significant deregulation of 14 cancer-related IPA pathways for which pathway directionality was known in HPDE *SPP1* knockout lines vs HPDE scr gRNA controls. 13/14 of these cancer-related pathways, including Pancreatic Adenocarcinoma Signaling, were in a direction suggestive that *SPP1* loss protects against tumor progression in human pancreatic duct cells. These findings are in agreement with published studies suggesting that *SPP1* loss ameliorates aggressiveness of pancreatic cancer cells (63, 64) and colon cancer cells (62, 65).

The requirement for Geminin in prevention of DNA re-replication initiation has been postulated to be when cells are stressed to divide quickly (66). We were unable to detect DNA damage with Geminin loss in homeostatic pancreatic duct cells, which may be due to the low proliferation rate of pancreatic duct cells and/or the presence of compensatory mechanisms with redundant function, such as ubiquitin-dependent degradation of Cdt1 at the time of replication licensing (67–70). Compensatory mechanisms are not sufficient to rescue the effects of Geminin loss in pancreatic duct cells in the context of CP, the result of which is accumulation of sustained DNA damage evident by γ-H2AX, but not ATR labeling. It has been previously reported that ATR is activated in Geminin-depleted colon cancer cell lines (71). Activation of the ATR-Chk1 pathway isn’t a major player in pancreatic duct cells in the setting of CP (72), suggesting different mechanisms participate in sensing Geminin depletion-induced DNA damage in different experimental systems and tissues.

## MATERIALS AND METHODS

### Preparation of pancreatic duct cells for single cell analysis

Pancreata from four 9 week old female C57BL/6J littermates (Jackson Labs, Stock 000664) were dissected, digested into single cells, and the DBA^+^ fraction obtained as previously described (12). Subsequently, live DBA^+^ cells were isolated for scRNA-seq by excluding propidium iodide (Thermo Fisher Scientific, P3566) positive single cells during FACS. scRNA-seq was performed by the Institute for Human Genetics Genomics Core Facility at University of California San Francisco (UCSF) using the 10X Genomics platform. Briefly, live, single, DBA^+^ pancreatic cells were loaded onto the microfluidic chip to generate single cell GEMs (Gel Bead-In EMulsions). Following cell lysis and unique barcode labeling, the cDNA library of 18,624 live pancreatic cells was generated using the Chromium Single Cell 3☐ GEM, Library & Gel Bead Kit v2 (10X Genomics). The cDNA library was sequenced on one lane using an Illumina HiSeq 4000.

### Single cell RNA-seq data processing

scRNA-seq data was generated on the 10X platform (10X Genomics, Pleasanton, CA) according to Single Cell 3’ protocol (v2 Chemistry) recommended by the manufacturer (73). The Cell Ranger software pipeline (version 2.1.1) was used to demultiplex cellular barcodes, map reads to the genome and transcriptome using the STAR aligner, and produce a matrix of gene counts versus cells. Doublets were filtered by excluding cells having RNA counts > 30000 and mitochondrial genes percentage > 10% in addition to using Scrublet (74). The R package Seurat (75) was used to process the unique molecular identifier (UMI) count matrix and to perform data normalization (gene expression measurements for each cell were normalized by total expression, and log-transformed), dimensionality reduction, clustering, ductal cell isolation, and differential expression analysis. We identified three clusters enriched in genes from 2 different cell types including: 1) acinar and T cell, 2) acinar cell and macrophage and 3) acinar cell and duct cell. Because our dataset doesn’t contain a population of acinar cells (they aren’t DBA^+^), doublet detector algorithms won’t remove acinar cell doublets from our dataset. Based on this reasoning, we removed these clusters containing a high threshold level of expression of acinar cell genes.

### Generation of *Geminin* conditional floxed allele

The general strategy to achieve Cre recombinase-mediated conditional gene ablation was to flank exons 3 and 4 of *Mus musculus Gmnn* by *loxP* sites (Figure S10A). The arms of homology for the targeting construct were amplified from BAC clone RP23-92G13 by PCR with high fidelity Taq polymerase. One primer contained a *loxP* site and a single SphI site which was used to verify the presence of the *loxP* site associated with it. Finally, the selectable cassette *CMV-hygro-TK* was incorporated into the targeting vector. The selectable marker itself was flanked by two additional *loxP* sites generating a targeting vector containing three *loxP* sites. Such a strategy allows the generation of ES cells with both a knockout allele and a conditional knockout allele after Cre mediated removal of the selection cassette *in vitro*. The targeting vector was sequenced to guarantee sequence fidelity of exons 3-4 and the proper unidirectional orientation of the three *loxP* sites. The complete left arm of homology was about 3200bp in length and the right arm of homology was 2100bp in length.

V6.5 ES cells were electroporated (25μF, 400V) with the three *loxP* sites-containing targeting construct, and hygromycin selection was performed to identify correctly targeted ES cells. Successfully targeted ES cells (*3loxP*) were identified with Southern blot (Figure S10B). These *3loxP* ES cells were then electroporated with a Cre-expressing plasmid and counter-selected with ganciclovir. ES cells that contained either one *loxP* or two *loxP* sites, respectively, were identified by Southern blot (Figure S10C). An ES cell clone was chosen that carried the conditional knockout allele (two *loxP* sites flanking exons 3 and 4) and was used for blastocyst injections to generate chimeric founder mice. *Gmnn*^f/f^ mice displayed normal litter sizes. For routine genotyping of *Gmnn^f^*^/f^ mice, the primers GCCTCGAACTCAGAAATCCA (primer A) and AACACAAAATTTGGCCTGCT (primer B) were used. To identify the deleted allele by PCR, primer C (TAGCCCGGACTACACAGAGG) can be used with primer A.

### Southern blot

For Southern blotting of genomic DNA, samples were digested with SphI or Bsu36I restriction enzymes for at least 4hrs and separated on an 0.8% agarose gel. The DNA was transferred to a Hybond-XL membrane (GE-Healthcare) in a custom transfer setup. Before assembly, the agarose gel was treated for 15min in depurination solution (21.5ml 37% HCl in 1L H_2_O), briefly rinsed in H_2_O and then soaked in denaturing solution (20g NaOH pellets, 87.6g NaCl in 1L H_2_O) for 30min. After transfer, the DNA was crosslinked to the membrane with UV light. The PCR amplified external Southern blot probes were labeled with ^32^P using the Prime-It II Random Primer Labeling kit from Stratagene. After hybridization of the probe and washing of the membrane, Kodak MS film was exposed to it and then developed.

### Mice

NSG mice from Jackson Labs (Stock 005557) were used. The transgenic mouse strain *Sox9– Cre^ERT^*^2^ was obtained from Jackson Labs (Stock 018829), and *Hnf1b–Cre^ERT^*^2^ has been previously described (53). Mice were maintained on a mixed genetic background. To induce Cre recombination, mice were injected with 6.7mg tamoxifen (Actavis, NDC 0591-2473-30) via oral gavage three different days over the course of a week at 7-9 weeks of age. Pancreatic duct ligations were performed as previously described (76). BrdU (Sigma, B9285-1G) injections were performed 24 hours and 4 hours prior to dissection. Mice were genotyped by PCR or Transnetyx. All animal studies were approved by the Institutional Animal Care and Use Committee at UCSF.

### Histology/immunostaining

Tissues were fixed in Z-Fix (Anatech Ltd., 174), processed according to a standard protocol, and embedded in Paraplast Plus embedding agent for histology, with DMSO (VWR 15159-464). For immunostaining, paraffin sections were deparaffinized, rehydrated, and antigen retrieval was performed, for all antibodies except BrdU, with Antigen Retrieval Citra (Biogenex, HK086-9K) using a heat-mediated microwave method. For immunostaining of BrdU, antigen retrieval was performed as previously described (77). For IHC, endogenous peroxidase activity was blocked by incubation with 3% hydrogen peroxide (Fisher Scientific, H325-100) following antigen retrieval. Primary antibodies were incubated overnight at 4°C. Secondary antibodies were used at 1:500 and incubated at room temperature for 1 hour (IHC) or 2 hours (IF). For IF, slides were mounted in ProLong Diamond Antifade Mountant with DAPI (ThermoFisher, P36962). For IHC, Vectastain Elite ABC kit (Vector Laboratories, PK-6100) and DAB Peroxidase (HRP) Substrate kit (Vector Laboratories, SK-4100) were used. Primary antibodies used in this study are listed in Table S2. Secondary antibodies used in this study were obtained from Life Technologies and Jackson Immunoresearch.

Immunostaining of cluster markers as well as the types of ducts within the ductal hierarchy tree were reviewed and classified by a board-certified pathologist. For expression analysis of selected markers in murine and human tissues, images shown are representative of at least 3 different donors or 9 week-old C57BL/6J mice. For quantification of BrdU, cleaved caspase 3, Geminin, Ki67, H2AX, and ATR, at least 60 cells from 3 different ducts were analyzed. For quantification of C-peptide/Pdx1/Nkx6.1 triple positive cells, at least 3 images containing an average of 188 cells were counted from 3 independent experiments per cell line. Normal human tissue used in this study was obtained from research consented human cadaver donors through UCSF’s Islet Production Core. Human pancreatic tissue specimens from five surgical resections from patients without pancreaticobiliary carcinoma or high grade pancreatic intraepithelial neoplasia were obtained. The pancreatic histologic section demonstrated chronic pancreatitis with loss of acinar parenchyma resulting in atrophic lobules along with variable fibrosis and chronic inflammation (most had no to sparse lymphocytic inflammation).

### Immunocytochemistry

Cells were grown on coverslips in 6 well plates and fixed at RT for 15 minutes with 4% paraformaldehyde. Cells were permeabilized with permeabilization solution (0.1% w/v Saponin, 5% w/v BSA in PBS−/−). The primary antibody was incubated in permeabilization solution at 4°C overnight. After washing off unbound primary antibody with PBS−/−, the secondary antibody was incubated in permeabilization solution for 1 hour at RT. After washing off unbound secondary antibody with PBS−/−, cells were mounted using ProLong Diamond Antifade Mountant with DAPI (ThermoFisher, P36962).

### RNA-seq

RNA was isolated using the RNeasy Mini Kit (Qiagen, 74106) as per manufacturer’s instructions. To obtain N=3 for the HPDE scr gRNA control, RNA was isolated on 3 different days of subsequent passages. A stranded mRNA library prep was prepared using PolyA capture and paired-end sequencing was performed by Novogene. 40 million reads were sequenced for each sample. Quality of raw FASTQ sequences was assessed using FASTQC. To process RNA-Seq libraries, adaptor sequences were trimmed using Cutadapt version 1.14 (requiring a length greater than 10 nt after trimming) and quality-filtered by requiring all bases to have a minimum score of 20 (-m 20 -q 20). Only reads that passed the quality or length threshold on both strands were considered for mapping. Reads were aligned to the human genome GRCh38 (hg38) with the STAR Aligner (version 020201). Ensembl reference annotation version 89 was used to define gene models for mapping quantification. Uniquely mapped reads for each gene model were produced using STAR parameter “--quantMode GeneCounts.” Differential expression analysis was performed in R using DESeq2 (v.1.16.0) with the default parameters, including the Cook’s distance treatment to remove outliers. The RNA-seq and scRNA-seq datasets were deposited to GEO (GEO accession #GSE159343).

### Cell culture assays

HPDE E6/E7 cells (78) were cultured in DMEM (Life Technologies 11995073), 10% FBS (Corning, 35011CV), 1X Penicillin: Streptomycin solution (Corning, 30-002-CI). For CellTiter-Glo Luminescent Cell Viability Assays (Promega, G7570), 1500 cells were seeded in quadruplicate per cell line in a 96 well plate in 150uL media, and luminescence was measured 5 days after plating cells. Values depicted for all cell culture experiments represent the average of at least 3 independent experiments.

For 3D proliferation assays, 1000 cells were seeded in triplicate per cell line in a drop of growth-factor reduced Matrigel (Corning, 356231) diluted 1:1 with complete HPDE media. After 4 days, images were taken of each well. For quantification, at least 40 individual spheroids per well were manually circled and the area determined using ROI tools from ImageJ (version 2.0.0). The average area of each well was normalized to the average from triplicate wells of the HPDE scr gRNA.

For carbonic anhydrase activity assays, cell lysates were prepared using standard protocols and cell lysis buffer (Cell Signaling Technologies, 9803S) containing 100 mM PMSF, 1X cOmplete Protease Inhibitor Cocktail (Roche, 11697498001), and 1X PhosSTOP (Sigma Aldrich, 4906845001). Carbonic anhydrase activity was measured using the Carbonic Anhydrase Activity Assay Kit (Biovision, K472-100). For normalization, equal amounts of protein (10ug) per sample were used in the assay. Protein concentration was determined using the Pierce BCA Protein Assay Kit (Thermo Fisher Scientific, 23225).

### Generation of stable knockout HPDE E6/E7 cell lines

For generation of stable knockouts, gRNAs were cloned into eSPCas-LentiCRISPR v2 (Genscript). gRNA sequences are included in Table S3. Each gRNA-containing plasmid was incorporated into lentivirus. HPDE E6/E7 cells were transduced with these lentiviruses, and cells expressing the gRNA-containing plasmid were selected for with puromycin. All cell culture experiments were performed using bulk transduced HPDE E6/E7 cells.

### Western blotting

Cell lysates were prepared using standard protocols and RIPA buffer (Thermo Fisher Scientific, 89901) containing 100 mM PMSF, 1X cOmplete Protease Inhibitor Cocktail (Roche, 11697498001), and 1X PhosSTOP (Sigma Aldrich, 4906845001). PVDF membranes were incubated with primary antibodies overnight at 4°C. After RT incubation with the appropriate HRP-conjugated secondary antibody for 1 hour, membranes were developed using SuperSignal West Pico PLUS Chemiluminescent Substrate (Thermo Scientific, 34580).

### Bioinformatics and statistical analysis

We used p ≤ 0.05 as a cutoff for DEG inclusion for IPA and IPA upstream regulator analysis. Due to low cell number and high similarity, some comparisons did not yield an acceptable number of statistically significant DEGs (≤25), and we used a relaxed p ≤ 0.1 as a cutoff for these in order to identify more targets. GSEA was performed on the identified DEGs with the GSEA software (version 3.0) in the pre-ranked mode, with the Reactome pathway dataset (version 7.2). For analysis of published single cell developmental biology datasets, GSM3140915 (E12.5 SW), GSM3140916 (E14.5 SW), GSM3140917 (E17.5 1 SW), and GSM3140918 (E17.5 2 SW) were used. The two E17.5 datasets were from the same animal and were merged. Ductal clusters were identified by expression of marker genes Sox9, Krt19, and Epcam. Data are presented as mean ± SEM and were analyzed in GraphPad Prism or Microsoft Office Excel. Statistical significance was assumed at a *P* value of ≤ 0.05. *P* values were calculated with the unpaired *t*-test. For interpretation of statistical results from unpaired *t*-test, * = *P* value ≤ 0.05, ** = *P* value ≤ 0.01, *** = *P* value ≤ 0.001, and **** = *P* value ≤ 0.0001. For all statistical analyses, outliers were identified and excluded using the Grubbs’ outlier test (alpha = 0.05) or ROUT (Q=10%).

## ACKNOWLEDGEMENTS

The authors thank Christina S. Chung, Debbie Ngow, and Melissa Campbell for their indispensable help with mouse colony maintenance, genotyping, and technical assistance. The authors wish to acknowledge Luc Baeyens and Michael S. German for helpful intellectual discussions. Graphical illustrations were developed in part by Jimmy Chen using BioRender.com. The *Gmnn* floxed mouse strain was generated in Rudolf Jaenisch’s laboratory. This work was supported by a Richard G. Klein Fellowship in Pancreas Development, Regeneration or Cancer (to A.M.H.). A.M.H. was additionally supported by F32 CA221114 and a Hirshberg Foundation for Pancreatic Cancer Research Seed Grant. A.A.R. and G.K.F. were supported by the UCSF Bakar ImmunoX Initiative. E.A.C. was supported by R01 CA222862. Work in M.H.’s laboratory was supported by R01 CA172045 and a grant from the Parker Institute for Cancer Immunotherapy (PICI). M.H. owns stocks/stock options in Viacyte, Encellin, Thymmune, EndoCrine, and Minutia. He also serves as SAB member to Thymmune and Encellin and is co-Founder, SAB and Board member for EndoCrine and Minutia.

