## Supplementary material for "Single cell transcriptome analysis defines heterogeneity of the murine pancreatic ductal tree": Table S1

### Slide 1
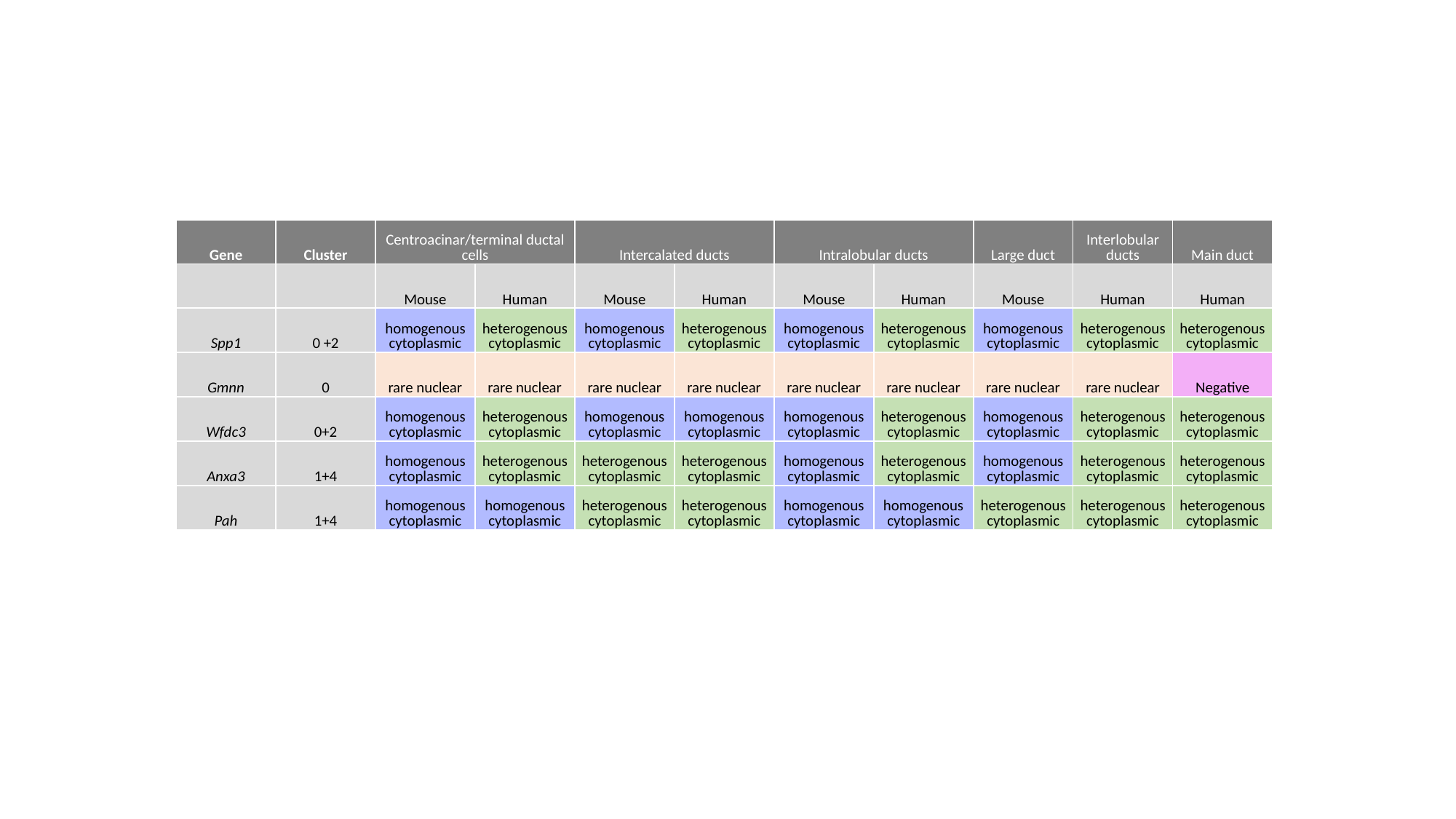

| Gene | Cluster | Centroacinar/terminal ductal cells | | Intercalated ducts | | Intralobular ducts | | Large duct | Interlobular ducts | Main duct |
| --- | --- | --- | --- | --- | --- | --- | --- | --- | --- | --- |
| | | Mouse | Human | Mouse | Human | Mouse | Human | Mouse | Human | Human |
| Spp1 | 0 +2 | homogenous cytoplasmic | heterogenous cytoplasmic | homogenous cytoplasmic | heterogenous cytoplasmic | homogenous cytoplasmic | heterogenous cytoplasmic | homogenous cytoplasmic | heterogenous cytoplasmic | heterogenous cytoplasmic |
| Gmnn | 0 | rare nuclear | rare nuclear | rare nuclear | rare nuclear | rare nuclear | rare nuclear | rare nuclear | rare nuclear | Negative |
| Wfdc3 | 0+2 | homogenous cytoplasmic | heterogenous cytoplasmic | homogenous cytoplasmic | homogenous cytoplasmic | homogenous cytoplasmic | heterogenous cytoplasmic | homogenous cytoplasmic | heterogenous cytoplasmic | heterogenous cytoplasmic |
| Anxa3 | 1+4 | homogenous cytoplasmic | heterogenous cytoplasmic | heterogenous cytoplasmic | heterogenous cytoplasmic | homogenous cytoplasmic | heterogenous cytoplasmic | homogenous cytoplasmic | heterogenous cytoplasmic | heterogenous cytoplasmic |
| Pah | 1+4 | homogenous cytoplasmic | homogenous cytoplasmic | heterogenous cytoplasmic | heterogenous cytoplasmic | homogenous cytoplasmic | homogenous cytoplasmic | heterogenous cytoplasmic | heterogenous cytoplasmic | heterogenous cytoplasmic |
