## Supplemental figure captions for "Single cell transcriptome analysis defines heterogeneity of the murine pancreatic ductal tree"

**Figure S1.** **Features of DBA^+^ (clusters 0-15) and ductal (clusters 0-5) cells.** A) The sorting strategy for live, DBA^+^ pancreatic cells is displayed. The graph on the left shows the cells plotted by forward and side scatter. In this graph, red cells are PI^+^, purple cells are DBA^+^PI^-^, and blue and green cells are doublets. The graph on the right shows the gating strategy used to sort DBA^+^PI^-^ pancreatic cells. B) The number of genes and transcripts for each cell in clusters 0-15 are shown. C) Co-immunofluorescence labeling identifies CK19^+^DBA^+^ and Collagen I^+^DBA^+^ copositive pancreatic cells and CD45^+^DBA^+^ copositive pancreatic lymph node cells. Pancreatic CD31^+^ cells are not DBA^+^. A yellow arrow points to a copositive cell. Scale bars are 50 µm. D) The plot shows whether a ductal cell is in cluster 0 or 8 (from the 0-15 cluster dataset) in the 0-5 cluster UMAP. E) This plot depicts the location of the ductal clusters 0-5 in the 0-15 cluster UMAP. F) Violin plots show the number of genes and transcripts in each cell for ductal clusters 0-5. G) Feature plots depict expression of genes normally enriched in pancreatic duct cells.

**Figure S2. IHC illustrates expression of markers in clusters 0 and 2 in the mouse and human ductal tree.** A) Geminin is expressed in rare ductal cells and acinar cells in the mouse and human pancreas. Yellow arrows point to Geminin positive cells. B) Osteopontin expression is observed in all duct cell types throughout the mouse and human ductal tree as well as in acinar cells. C) Wfdc3 is expressed in all duct cell types in the mouse and human pancreas and in acinar cells. Red arrows point to the indicated duct type. Scale bars are 40 µm.

**Figure S3. IHC and IF depict expression of markers in clusters 1, 3, 4, and 5 in mouse and human pancreas duct cells.** A) Annexin A3 expression is observed in all ductal cell types in the mouse and human pancreas. The left mouse IHC image under Large duct – Interlobular/Main duct shows Annexin A3 cytoplasmic expression in pancreatobiliary cells and cells within the Ampulla of Vater. Scale bars are 40 µm. B) Pah is expressed in all duct types throughout the mouse and human ductal tree as well as in acinar cells. Scale bars are 40 µm. C) Yellow arrows point to heterogeneous expression of ductal markers Cftr, Annexin A3, and CK19 in human pancreatic duct cells. Scale bars are 20 µm. D) Proliferating and acetylated alpha tubulin positive duct cells are observed in the intrapancreatic bile duct, peribiliary glands, and pancreatobiliary cells in mouse and human. Scale bars are 50 µm.

**Figure S4. Characteristics of intrapancreatic bile duct and pancreatobiliary cells.** A) Cxcl5, another marker of *Dmbt1*^+^*Ly6d*^+^ cells, positive cells are located in murine and human intrapancreatic bile duct cells, peribiliary glands, and pancreatobiliary cells. Yellow arrows point to ductal cells displaying upregulated Cxcl5. Scale bars are 50 µm. B) Stacked violin plots show expression of 5 genes which are upregulated in clusters 3 and 4. C) The dot plot shows the top 20 DEGs ranked by fold change when comparing clusters 3 vs 4.

**Figure S5. Alignment to an adult murine hepatic biliary epithelial cell dataset.** A) UMAP showing alignment of adult murine hepatic BECs (blue) to our murine intrapancreatic bile duct cells (red) and pancreatobiliary cells (green). B) Clustering of merged datasets defines 5 clusters. C) Intrapancreatic bile duct cells in DBA^+^ duct cluster 3 are primarily located within the merged clusters 0 and 1, and pancreatobiliary cells in DBA^+^ duct cluster 4 are primarily located within the merged clusters 1 and 2. The heatmap shows the percent of cells from our clusters 3 and 4 within each of the merged clusters 0-4. D) Feature plots depict the 75^th^ percentile and higher of cells expressing the published gene signatures of hepatic BEC subpopulation A and B respectively. E) Cells in clusters 1, 2, and 4 have the strongest enrichment for subpopulation A genes, while cells in clusters 0, 1, and 3 have the strongest enrichment for subpopulation B genes in the merged dataset. F) Dual violin plots show expression of the ductal marker *Sox9* and the Yap targets *Cyr61*, *Ankrd1*, and *Gadd45b* in the merged clusters. G) Dot plot shows expression of hepatic BEC subpopulation A and B genes, analyzed by t-SNE, in Figure 1D of Pepe-Mooney *et al.* (2019) in our murine pancreas DBA^+^ duct clusters 0-5.

**Figure S6. Analysis of pancreas duct cells during development.** A, E, I) UMAPs depict ductal clusters evident at E12.5, E14.5, and E17.5, respectively. B, F, J) Cluster dendrograms created using dims (used to define the cluster) shows the Euclidean relationships between clusters at E12.5, E14.5, and E17.5, respectively. C, G, K) Dot plots show expression of the top 5 genes defining adult C57BL/6J duct subpopulations in developmental biology samples at E12.5, E14.5, E17.5, respectively. M-N) Feature plots of genes that characterize subpopulations of mouse and human duct cells in Baron *et al.* (2016). O) Feature plot showing *Fth1* expression, which typifies a human pancreas duct subpopulation in Grün *et al*. (2016).

**Figure S7. Comparison of DBA^+^ lectin sorted mouse pancreas duct subpopulations to ALK3^+^ human pancreas duct subpopulations.** A-E) Aggregated expression of control feature sets shown in panel A were subtracted from the average expression levels of DEGs for each cluster 0-5 on a single cell level to determine the AddModuleScore comparing each DBA^+^ pancreas ductal cluster to ALK3^+^ human pancreas clusters. Panel D shows the number of DEGs in murine DBA^+^ pancreas duct clusters 0-5 that have a human homolog and could be used in this comparison. F) *Bmpr1a* is expressed in a subset of murine DBA^+^ pancreas duct cells. G) Stacked violin plots depict expression of centroacinar/terminal ductal cell markers *Hes1*, *Aldh1b1*, *Aldh1a1*, and *Aldh1a7* in DBA^+^ pancreas duct clusters 0-5. H) Immunostaining identifies ducto-acinar cells in murine pancreas. Yellow arrows point to Cpa1 or α-amylase positive murine ductal cells. Similar to other murine ductal cell markers (Figure S1G), DBA lectin also shows heterogenous expression in murine pancreatic duct cells. Blue arrows point to a DBA lectin negative duct cell. The scale bar is 50um.

**Figure S8. RaceID3 clusters and Monocle 3 analysis.** A) The location of DBA^+^ duct cluster 0, 1, 2, and 4 cells in RaceID3 clusters are depicted in Seurat space. Clusters 3 and 5 were not included in this analysis and all clusters with <10 cells were also removed. All removed cells are depicted in the N/A square. B) RaceID3 cluster 10 cells, which have the highest StemID2 score, are depicted in the Monocle 3 UMAP. C) The Monocle 3 UMAP and trajectory are shown when DBA^+^ duct clusters 0-5 are all included in the analysis. D) A violin plot showing the distribution of cells along the relative pseudotime axis, split by DBA^+^ duct clusters, is shown. Cluster 4 appears to be the most differentiated from the inferred root, cluster 0.

**Figure S9. Characterization of DBA^+^ murine ductal markers.** A) Representative images for 3D proliferation assays are shown. Scale bars are 100 µm. B-E) GSEA enrichments plots for select pathways are depicted.

**Figure S10.** **Generation of 2loxP and 1loxP heterozygous ES cells for mouse *Geminin.*** A) The targeting strategy to generate a conditional KO allele for *Geminin* in ES cells is shown. The exact distance between individual exons and their relative sizes is not shown. ES cells heterozygous for the 3loxP allele were obtained through homologous recombination. A Cre recombinase was used to generate ES cells harboring either the 2loxP allele or the 1loxP allele *in vitro.* A SphI restriction site was introduced with the leftmost *loxP* site to allow screening for its presence by Southern blot analysis. B) The 5’ probe was used in conjunction with a SphI digest. Besides the wild-type allele, a fragment of about 2.5kb is expected for the 3loxP allele as indicated. Clone 17 and clone H18 tested positive. C) Clone H18 was chosen for Cre treatment *in vitro*. Using a Bsu36I digest and the 5’ probe, the 2loxP allele displays a single fragment of the same size as the wild-type allele whereas the 1loxP allele produces a smaller fragment. Bsu36I restriction sites are omitted in the schematic shown in A) for clarity. The 3’ probe clearly distinguishes between wild-type, 1loxP, 2loxP, and 3loxP alleles with a SphI digest. Clone 53 was identified as an ES cell clone heterozygous for the conditional 2loxP allele and used for blastocyst injection.

**Figure S11. Histology of transgenic PDL models.** A-B) Representative H&E images of PDL transgenic models at Day 7 and Day 30 are shown. Scale bars are 40 µm.

**Figure S12. Geminin loss causes a transient proliferation response in *Sox9cre^Tg/wt^*; *Geminin^f/f^* mice.** A) Representative IF images of the quantification shown in Figure 7D are depicted. Scale bar is 20 µm. B-C) IF images and quantification of BrdU positive pancreatic ductal cells is shown at Day 7 in the PDL transgenic models. Scale bars are 50 µm. D-E) IF images and quantification of BrdU positive pancreatic ductal cells is shown at Day 30 in the *Sox9cre^Tg/wt^*; *Geminin^f/f^* and control PDL models. Scale bars are 50 µm.

**Table S1. Expression scoring of markers of subpopulations of pancreatic duct cells.** This table depicts a summary of expression scoring of selected markers for subpopulations of pancreatic duct cells in mouse and human tissue. Homogeneous refers to an observed uniform expression level and pattern within a particular ductal cell type. Heterogeneous means that either the observed expression level or pattern varies among cells within a particular ductal cell type.

**Table S2. The list of antibodies used in this study.**

**Table S3. The list of gRNA sequences used in this study.**

**Dataset S1. Number of cells and average number of genes and transcripts in all Seurat clustering analyses.**

**Dataset S2. DEGs in various comparisons.**

**Dataset S3. IPA results.**

**Dataset S4. IPA Upstream Regulator analysis results.**

**Dataset S5. Monocle 3 module analysis.**
